# Electrocatalytic on-site oxygenation for transplanted cell-based-therapies

**DOI:** 10.1101/2023.06.05.543794

**Authors:** Inkyu Lee, Abhijith Surendran, Samantha Fleury, Ian Gimino, Alexander Curtiss, Cody Fell, Daniel Shiwarski, Omar El-Refy, Blaine Rothrock, Seonghan Jo, Tim Schwartzkopff, Abijeet Singh Mehta, Sharon John, Xudong Ji, Georgios Nikiforidis, Adam Feinberg, Josiah Hester, Douglas J. Weber, Omid Veiseh, Jonathan Rivnay, Tzahi Cohen- Karni

## Abstract

Implantable cell therapies and tissue transplants require sufficient oxygen supply to function and are limited by a delay or lack of vascularization from the transplant host^1, 2^. Exogenous oxygen production can support cells and tissues, such as pancreatic islets and engineered therapeutic cells. Previous oxygenation strategies have targeted gas circulation or decomposition of solid peroxides. These strategies however require bulky implants, transcutaneous supply lines, and are limited in their total oxygen production or regulation^3, 4^. Readily integrated and controlled production of oxygen has eluded cell therapy devices. Here, we show an electrocatalytic approach that enables bioelectronic control of oxygen generation in complex cellular environments to sustain engineered cell viability and therapy production under hypoxic stress and at high cell densities. Nanostructured sputtered iridium oxide serves as an ideal catalyst for oxygen evolution reaction (OER) at neutral pH. It enables a lower OER onset and shows selective oxygen production without evolution of toxic side products over a 300 mV window of operation. This electrocatalytic on site oxygenator (ecO_2_) can sustain high cell loadings (>60k cells/mm^3^) in hypoxic conditions in vitro and in vivo. Our results demonstrate that exogenous oxygen production devices can be readily integrated into bioelectronic platforms and enable high cell loadings in smaller device footprints with broad applicability.

## Main

The transplantation of therapeutic allogeneic or xenogeneic cells, within semipermeable devices as a living pharmacy has the potential to treat a range of diseases such as endocrine disorders, autoimmune syndromes, cancers, and neurological degeneration^5–9^. Cell-based therapeutics translation to humans requires high cell densities^2^ to enable miniaturized devices of therapeutic value. However, maintaining the potency of such densely packed therapeutic cells for extended duration is challenging due to a number of factors such as immune response from the host and inadequate availability of nutrients and dissolved oxygen^8, 10^. While immunoisolation and cell protection using encapsulating size-exclusive membranes or engineered biomaterials can partially address immunoreactivity, nutrient and oxygen insufficiency presents a critical challenge. Oxygen has been regarded as the limiting factor supporting cell viability and potency^1^. Due to the oxygen mass diffusion limit, in a native tissue each cell is within *ca*. 100 µm from a blood capillary to allow adequate oxygen supply^1^. Transplanted exogenous cells or tissue require neovascularization or supplemental oxygenation^11^. Delay in vascularization limits the success of the transplantation as well as the transplanted cells’ function. Oxygen deficiency in the transplanted cells is caused by (1) insufficient oxygen tension at implantation site, (2) innate large oxygen consumption of cells (i.e., metabolic demand), (3) high cell density, (4) additional barriers to oxygen diffusion (i.e., membranes, or formation of encapsulating fibrotic tissue).

To address the hypoxic stress on transplanted cells, various strategies have been investigated to enhance exogenous oxygen delivery. Two major strategies employed to mitigate oxygen deficiency may be classified into active and passive methods. Active methods involve oxygen release through an externally controllable mechanism, e.g., delivery of gaseous oxygen to the transplanted cells (i.e., islet cells)^3^. Passive methods rely on gradual release of oxygen through unregulated or self-regulated mechanisms, e.g., engineered platforms to increase the oxygen exchange with the implantation environment^12^ or release of oxygen from metal peroxides^4, 13, 14^. Though these approaches are able to support transplanted cells they are limited in control of oxygen release, lifetime of available oxygen supply and limited supported cells’ density, e.g., pancreatic islet (10k cells/mm^3^)^15^ and engineered cells (6k cells/mm^3^)^16^.

Electrochemical water electrolysis for oxygen production is a promising approach for providing oxygen to cells ^17, 18^. However, its demonstration *in vivo* has been limited due to improper materials selection geared toward efficient water splitting, the use of bulky and complex electronics for splitting water and limited power budget. Although electrocatalytic water splitting is widely accepted in renewable energy research, e.g., fuel cells, its application in tissue engineering has been limited due to the sluggish nature of water splitting in neutral environments^19^. The pH-dependent nature of water’s redox reactions and its inherent thermodynamic stability further hinder electrochemical water decomposition in physiological environments, as they lack the necessary active species like H^+^ and OH^-^ ^20^. Additionally, neutral pH necessitates a higher overpotential, leading to increased power consumption and the potential risk of generating toxic byproducts such as chlorine through chloride oxidation (*E*_OX_(Cl^-^) = 1.771 V vs. RHE). Consequently, it is crucial to employ electrocatalysts that can effectively lower the energetic barrier while being biocompatible and highly selective for oxygen evolution reaction (OER) at neutral pH.

Iridium oxide is a benchmark electrocatalyst for OER due to its exceptional catalytic efficiency and stability^21, 22^. Unlike other electrocatalytic materials, iridium oxide retains its robust catalytic activity in acidic environments where water molecules are directly involved in the OER^19, 22–25^. This makes it suitable as an electrocatalyst in physiological environments, as the electrochemistry of OER is fundamentally similar in neutral and acidic conditions^20^. Its biocompatibility has been validated by extensive research outcomes in the electrophysiology community^26–30^. Iridium oxide films can be produced with nanoscopic morphologies, which allow device miniaturization and enlarged electrochemical surface area readily available for electrocatalysis^27, 29, 30^.

Here we report a highly controlled, on demand electrocatalytic on-site oxygenator (ecO_2_) platform designed to safely support implanted therapeutic cells. Capitalizing on its unique catalytic properties, stability, patternability and biocompatibility, we employed a nanostructured sputtered iridium oxide film (SIROF) to enhance the kinetics of OER and reduce the energetic cost to produce oxygen. Detailed engineering of the ecO_2_ geometry resulted with precise control over the distribution of generated oxygen. In vitro experiments assessed the platform’s ability to sustain cell viability while maintaining therapeutic peptide secretion in high-cell density (60k cells/mm^3^) under hypoxic conditions (1% O_2_) up-to 3 weeks. ecO_2_ effectiveness was subcutaneously tested in vivo (rat) for 10 days, showing its promise as a tool for cell-based therapeutic interventions, and paving the way for biomedical engineering applications, e.g., transplanted cells to treat diabetes^31^ or cancer^32^.

### De Novo design of ecO_2_

The ecO_2_ platform comprises a specialized chamber for housing high-density therapeutic cells and an integrated oxygenation system to provide oxygen support, implanted at readily accessible and clinically relevant locations such as subcutaneous and intraperitoneal sites (**Figure 1a**). Oxygen generation is achieved through electrolysis of water and precisely regulated using a battery powered and wirelessly controllable electronic system, although a fully battery-free wireless power technology can be readily implemented with ecO_2_. As a key material for oxygen production, sputtered iridium oxide (SIROF), which is a highly active and biocompatible electrocatalyst for water oxidation, was adopted in our system. To ensure uniform oxygen delivery toward implanted cells, we employed finite element analysis (FEA) to estimate the diffusional behaviors of the generated oxygen with various array geometries (**Figure 1b**). Given the symmetry of the designed electrode, 2-dimensional oxygen profiles were studied for cross-section of the electrode (**Supplementary Figure 1** and **Supplementary Note 1**). Based on simulation studies demonstrating a uniform oxygen distribution profile, we utilized an array composed of point sources (**Figure 1c**). The fabricated electrode configuration consisted of a central SIROF anode and a surrounding platinum cathode, resulting in highly distributed oxygen profiles. Fabrication methods and procedures are illustrated in **Materials and Methods** and **Supplementary Figure 2**. Whereas SIROF anode possesses smaller geometric surface area in comparison to Pt cathode, nanostructured SIROF enhances catalytic activities of iridium oxide toward oxygen evolution reaction, thus contributing to an enlarged electrochemical surface area. We maximized the surface area of the cathode where hydrogen is evolved to minimize current density and prevent bubble formation resulting from the cathodic reaction, which occurs due to the inferior solubility of hydrogen in water^18^. The synthesized catalysts for ecO_2_ exhibited nanoporous structure, which is advantageous for triggering electrocatalytic oxygen evolution reaction at lower potential (**Figure 1d**), thereby reducing power requirements for the electronics as well as preventing electrochemical side reactions which can occur upon high potential application. Surface analysis evidenced the presence of rutile IrO_2_ interspersed with amorphous regions (**Supplementary Figure 3**), as well as the existence of electrocatalytically active Ir^4+^ and oxygen vacancies for OER (**Supplementary Table 3** and **4,** and **Supplementary Figure 4**). The combination of highly active amorphous iridium oxide and stable crystalline IrO_2_ enabled the support of implantable cell therapies through chronic oxygen delivery via electrochemical water splitting^33^, recapitulating the critical features necessary for on-demand oxygen production.

**Figure 1.**
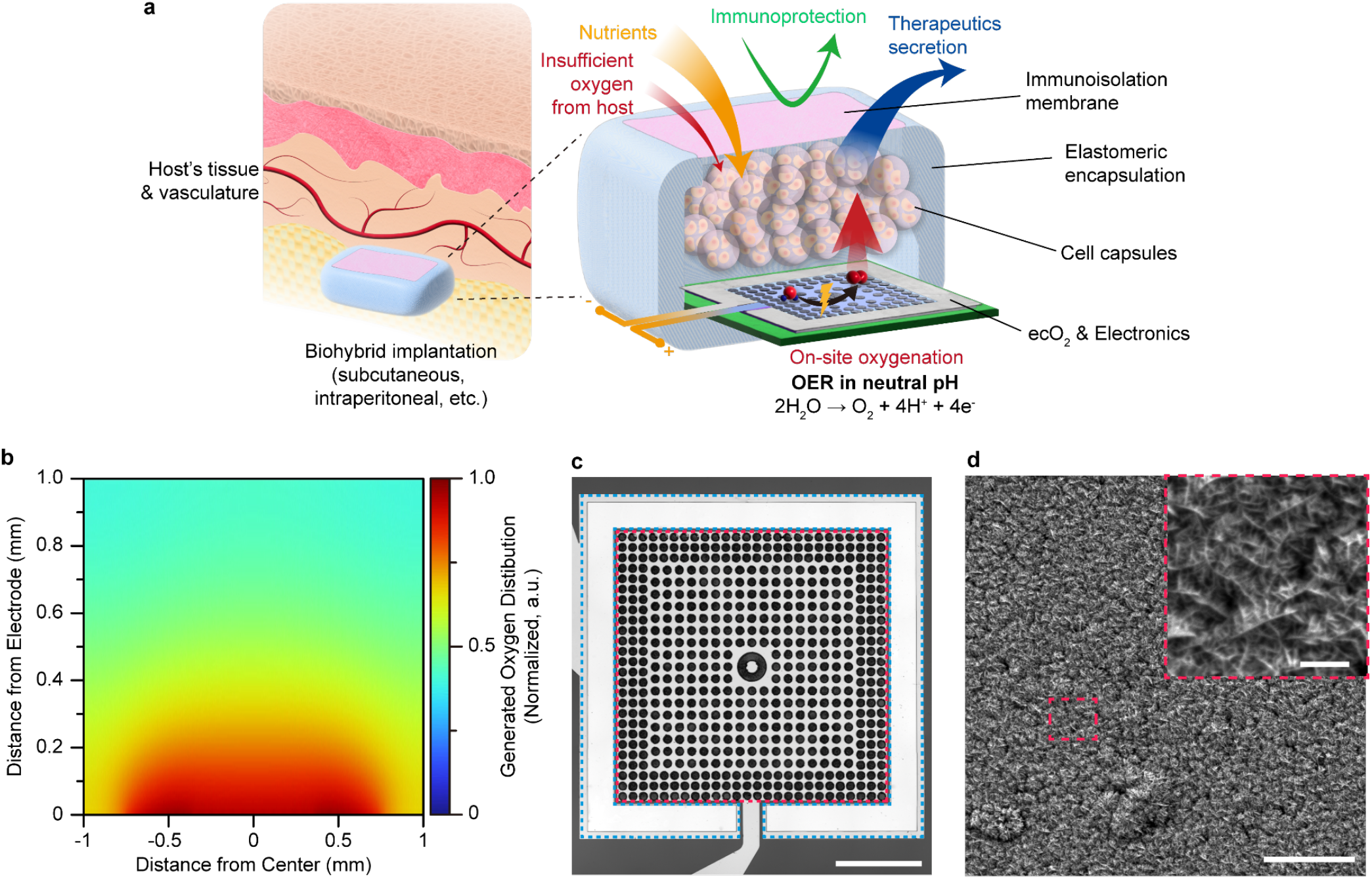
Electrocatalytic on-site oxygenation (ecO_2_) for implantable cell therapies platform. (a) A schematic illustration of ecO_2_. (b) A representative optical image of fabricated ecO_2_ device; blue dashed outline: Pt counter electrode; red dashed outline: working electrode with micropatterned circular sputtered iridium oxide (SIROF) catalytic array; scale bar: 500 µm (c) Oxygen distribution profiles from the designed catalytic arrays. Oxygen levels were normalized with the value at the bottom of the chamber (y=0 mm). (d) A representative scanning electron microscopy (SEM) image of SIROF; scale bar: 5 μm; inset: 500 nm.

### Effective, stable, and safe on-site oxygen delivery

We benchmarked the activity and stability of the SIROF-based catalyst electrode against platinum electrode in 1X PBS electrolyte (pH=7.4). Linear sweep voltammetry (LSV) scan in anodic direction revealed the characteristic activation signal of iridium oxide in OER at approximately 1.6 V vs. RHE^34, 35^. The SIROF catalyst demonstrated significantly lower overpotential of 395±18 mV compared to Pt electrode’s 565±10 mV (**Figure 1a**) representing reduced energetic cost to produce oxygen with catalytic activity of SIROF. Higher current observed for SIROF at the same applied potential implied its ability to achieve power-efficient electrocatalytic oxygen generation in bioelectronic applications. The stability of SIROF was examined under continuous chronoamperometric operation with a 2-electrode setup at 1.7 V over 14 days. To monitor the change in catalytic activities, LSV curves in 3-electrode and 2-electrode setup were collected every 24 hours. The onset of oxygen evolution for SIROF remained largely unchanged during the 14-day experiment, indicating that the applied catalyst did not experience significant degradation under continuous electrical stress (**Figure 2b)**. Additionally, the negligible onset change observed in 2-electrode settings indicated the overall stability of ecO_2_ over 14 days of oxygen evolution (**Supplementary Figure 5**).

**Figure 2.**
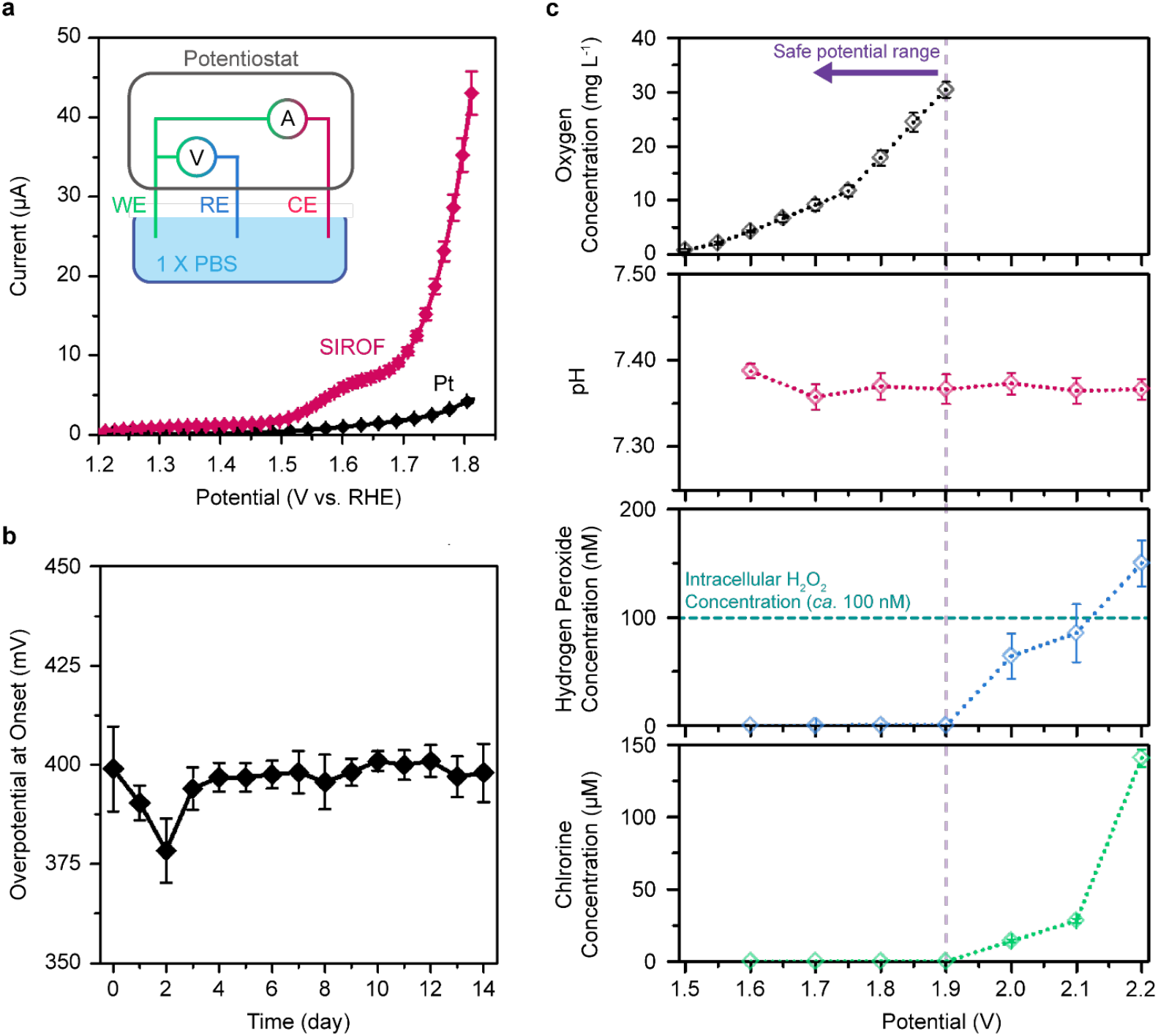
ecO_2_ is an effective, stable and safe oxygen generation platform. (a) Linear sweep voltammetry (LSV) measurement in anodic direction of pristine Pt (*n*=8, gray) and sputtered iridium oxide (SIROF) electrocatalysts (*n*=8, red) ; inset – An illustration of the 3-electrode electrochemical characterization setup ; WE - working electrode, catalyst array; RE - reference electrode, 1 M Ag/AgCl; CE - counter electrode, Pt mesh. (b) The stability of SIROF as an electrocatalyst, presented with overpotential change over 14 days at onset (3-electrode setup, *n*=5). (c) Oxygen generation at different applied potential and its side reactions; oxygen concentration - black; pH - red; hydrogen peroxide - blue; chlorine - green. Oxygen was produced with 30-sec pulse duration at each potential, while side reactions were detected after 24-hour chronoamperometry at each potential.

ecO_2_ can evolve oxygen in a highly controlled manner, without producing deleterious side products. Oxygen concentration was measured during potential pulsing (duration: 30 sec). ecO_2_ was able to produce oxygen at levels exceeding atmospheric levels (21 %, 8.238 mg L^-1^ at 25 ℃) when applying potentials greater than 1.6 V. Furthermore, bioelectronic control allows for modulation of oxygen tensions in a precise manner, by applying different potentials (**Supplementary Figure 6a** and **6b**) or changing the duty cycling (**Supplementary Figure 6c** and **6d**). Lower duty cycles provide an added benefit of extended device lifetime by curtailing power consumption and electrochemical stress on the catalytic arrays. While oxygen generation is validated, selectivity of oxygen production must also be confirmed in complex physiological environments. To this end, we monitored possible side-reactions such as chloride oxidation, peroxide formation, and pH change during 24-hour continuous oxygen generation. Since OER in neutral pH results in protons as a byproduct (2H_2_O → O_2_ + 4H^+^ + 4e^-^), local pH change may occur. However, pH was maintained within the buffer capacity of 1X PBS, which adequately mimics the buffering capacity of body fluids^36^. Measurements of evolved hydrogen peroxide showed detectable levels at potentials above 1.9 V. Even so, the concentration of generated hydrogen peroxide remained below the intracellular level of *ca*. 100 nM up to 2.2 V^37^. Finally, chlorine, which comes from chloride oxidation (2Cl^-^ → Cl_2_ + 2e^-^), was not detected below 1.9 V. Since chloride is one of the most abundant ions in biofluids, its oxidation was the most concerning side reaction. These combined results establish a wide operation window for safe and selective oxygen evolution of ∼300 mV, where OER onset occurs at 1.6 V, and harmful side reactions begin at 1.9 V (**Figure 3c**). This window may be further expanded to higher voltages with implementation of a selective coating or membrane that blocks reactants (i.e. Cl^-^ ions) from reaching the working electrode. It is this window of operation that enables the highly tunable on-site production of oxygen to support high cell densities of non-native cells.

**Figure 3.**
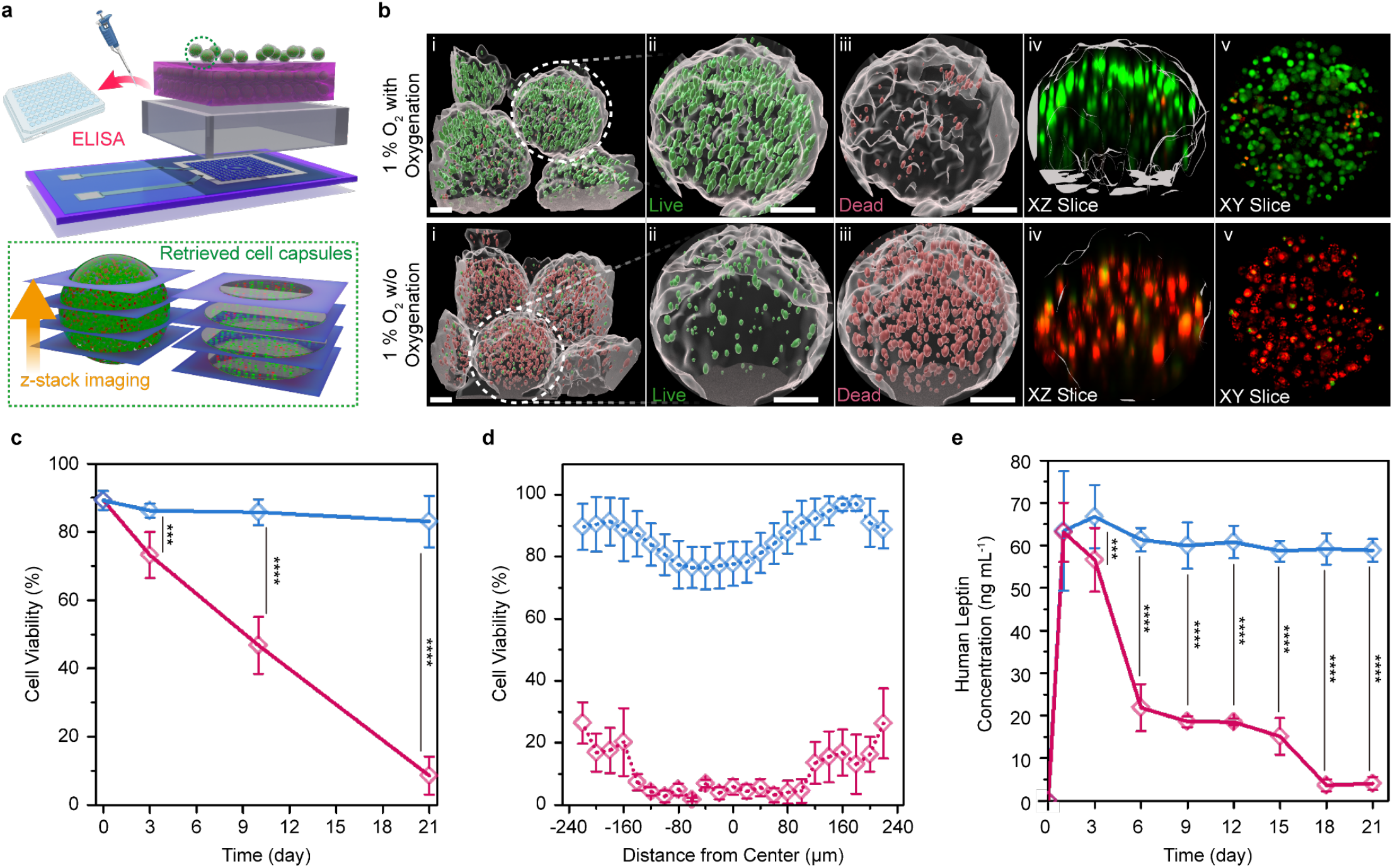
ecO_2_ supports cell capsules *in vitro* for 21 days. (a) A schematic illustration of the in vitro oxygenation assay and analysis. (b) A representative set of 3D-rendered immunofluorescence images after 21-day in vitro oxygenation; i - a representative 3D-rendered z-stacked images, ii - Expanded view of the marked white dashed circle, live cells, iii - dead cells, iv - cross-sectional view of the same capsule capsule in xz plane, v - cross-sectional view of the same capsule in a xy plane. All scale bars are 100 µm (c) Cellular viability as a function of time, (d) Spatial distribution of cell viability in cell laden alginate capsules. (e) Peptide production as a function of *in vitro* culturing time. Blue corresponds to 1 % oxygen with ecO_2_ (oxygenation in hypoxia, experiment, *n*=4); red corresponds to 1 % oxygen without ecO_2_ (hypoxic incubator, negative control, *n*=4); ***: *p* < 0.001; ****: *p* < 0.0001.

### ecO_2_ safely sustains high cell density *in vitro*

Supplemental oxygenation is required to sustain the metabolic needs of cells when packed at high densities. When there is an imbalance of O_2_ availability to sustain dense cell needs, encapsulated cells undergo apoptosis^15^. To test this hypothesis, we loaded ARPE-19 cells in spherical capsules (diameter of 415±31 μm) at 60k cells/mm^3^ density and exposed cultured in vitro for 21 days with and without O_2_ (For additional information see Materials and Methods). ARPE-19 cells were chosen as a model chassis for investigation because of their clinical relevance to a wide range of cell-based therapeutic products for oncology^7, 32^ (NCT05538624), eye disorders (NCT04577300), Blood disorders (NCT04541628), enzyme replacements (NCT05665036), and Alzheimer’s^38^. This cell line is widely used in the clinic because it is non-tumorigenic^39^, displays contact-inhibited growth characteristics^40^, amenable to genetic modification^39^, and in human trials, it has been shown to be safe^38, 41, 42^. By using a non-dividing cell line, we are able to assess potency losses because of cell death reliability owing to the fact that in our device, the transplanted ARPE-19 cells would not be mitotically active. By contrast, previous studies^8, 43^ with encapsulated cell therapy devices focused on non-clinically translatable cancer cell lines, i.e., HEK 293 cell chassis, which have the mitotic capacity in vivo and new cell replication can compensate for cell death from hypoxia.

High density cell clusters encapsulated in alginate capsules were adopted as an in vitro model to simulate high cell density transplantation. Approximately 100 capsules were added in each catalytic array. While nutrients other than oxygen were supported by periodic media exchange, it was assumed that oxygen supplementation was exclusively dependent on ecO_2_ oxygen production, because all cell culture media was deoxygenated under 1 % oxygen. Evaluated at different time points using fluorescence confocal microscopy (**Figure 3a**), live/dead assay of the oxygenated cells showed improved cell viability of 83.1±7.5 % after 21-day incubation in hypoxia, while hypoxic incubation without oxygenation (negative control) showed massive cell death (viability: 8.6±5.6 %) (**Figure 3b**). The obvious contrast in 3D-rendered cell capsule images confirmed that oxygen is indeed the limiting factor of high cell densities. While progressive decline of cell viability in negative control was attributed to hypoxic stress, only 7.5 % of live cells were dead after 3-week culture compared to the initial stage of culture (89.8±2.8 % at day 0) (**Figure 3c**). Notably, SIROF catalysts that underwent a continuous 21 days 100 % duty cycle exhibited nanoporous morphology with apparent degradation of the dendritic features (**Supplementary Figure 7**). This degradation was also observed in SIROF used in continuous stimulation of neural electrodes^42^. Interestingly, such significant oxygen demand in highly dense cell capsules led to hypoxic cell death even under normal oxygen level (20 %, positive control) (**Supplementary Figure 12** and **13**), confirming that on-site oxygen production beyond atmospheric oxygen level is crucial to address oxygen deficiency for dense or large cell constructs. Cell viability distribution along the z-axis showed no hypoxic core formation for the cell-loaded capsules that were oxygenated with ecO_2_ (**Figure 3d**). Though viabilities were slightly declined in the vicinity of the center, slice images in xy-plane did not exhibit necrotic core formation in the center at different z-positions (**Supplementary Figure 14**). In contrast, the cells from the negative control were only viable near the surface of the capsules, and live cells were essentially absent in the capsule core (**Supplementary Figure 15**). ecO_2_ demonstrated that hypoxic core formation in densely packed cell clusters is mitigated with on-site oxygenation, overcoming the limitation of oxygen diffusion from native vasculature.

While the viability of cells was improved by electrochemical oxygenation, the production of a peptide (or other transcriptional biomolecule) directly indicates maintenance of cellular potency toward applications in on-site therapy production. In fact, cell functionality is more sensitive to oxygen tensions because the dysfunction emerged at higher levels of oxygen than hypoxia-induced cell death^44^. The ARPE-19 cells herein were engineered to produce and secrete leptin, a therapeutic hormone^45^ used for applications in obesity, endocrine disorders, and regulation of circadian rhythm. Leptin levels in vitro were quantified via human leptin enzyme-linked immunosorbent assay (ELISA) over a 24h production period. Noticeably, the concentration of secreted leptin from the engineered ARPE-19 cells represented a similar tendency to viability, as depicted in **Figure 3e**. While the amount of the produced leptin declined over time in negative control, cells in the ecO_2_ oxygenation condition showed consistent levels of leptin production and release over the 3 weeks in vitro at high cell loadings.

### ecO_2_ supports implanted cells *in vivo*

Oxygen tension changes in different tissues, e.g., *ca*. 50, 40 and 30 mmHg O_2_ in the kidney, pancreas, and general intraperitoneal space respectively^46^. To test the efficacy of the ecO_2_ in vivo we chose to implant the platform subcutaneously where the oxygen tension is lower (39.0±6.3 mmHg O_2_). We demonstrated oxygen delivery to implanted cells in a rodent model (for additional information see Materials and Methods). While medical grade PDMS served as impermeable housing for electronics, a microporous polycarbonate membrane was added at the top of the cell compartment to provide selective mass transport for immunoisolation (**Supplementary Figure 18**)^8^. The implanted ecO_2_ consisted of a cell capsules compartment supported by 8 catalytic arrays for *ca*. 800 alginate cell capsules (60k cells/mm^3^). The ecO_2_ was implanted at the abdominal skin pocket and operated by a remote-controllable potentiostat circuit secured on rat’s back which was connected to the cell compartment via a flexible tether (**Figure 4a, Supplementary Figure 18** and **19**). The devices were retrieved and analyzed to evaluate cell viability after 10 days. Notably, there was no significant inflammation and foreign body responses, which implied successful immunoisolation of implanted cells from the host (**Supplementary Figure 20**). Capsules with ecO_2_ support showed significantly higher cell viability (73.7%) than capsules without ecO_2_ support (26.6%) (**Figure 4b**). The relatively lower viability compared to 10-day in vitro oxygenation implies that the volume of media per cell (in vitro: 2.5 μL per capsule, in vivo: 0.27 μL per capsule) and media exchange may affect the viability of the cells. Nonetheless, as depicted in **Figure 4c**, significantly improved viability of the oxygenated cells highlighted the effectiveness of ecO_2_ to support implantable cell therapies.

**Figure 4.**
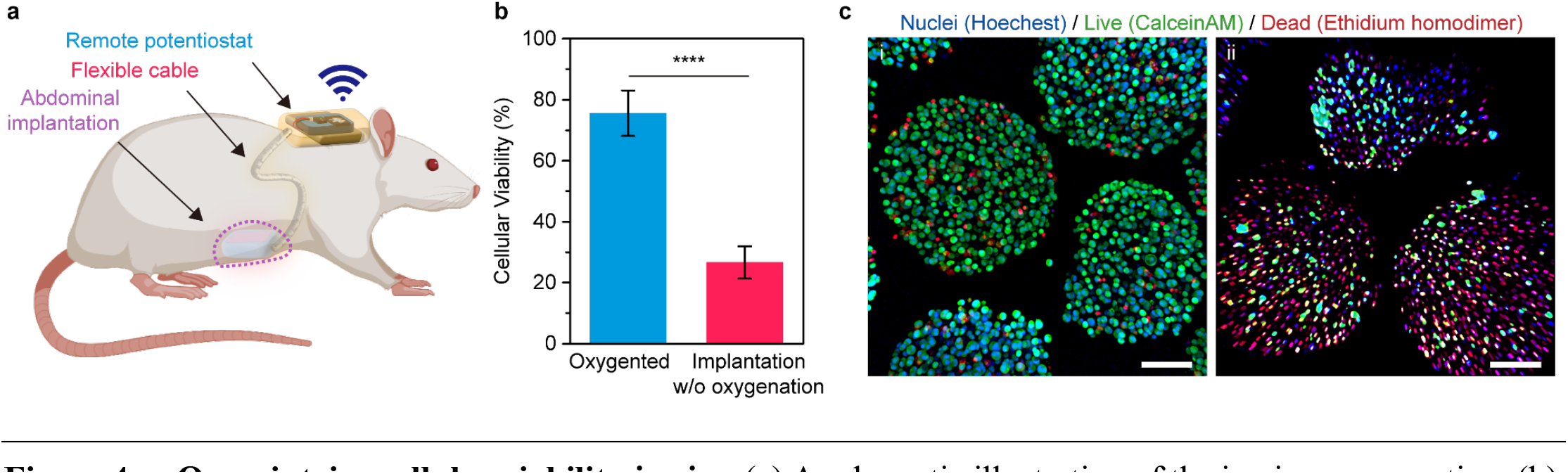
ecO_2_ maintains cellular viability *in vitro*. (a) A schematic illustration of the in vivo oxygenation. (b) The cell viability after 10-day implantation in rat with (blue) or without oxygenation (red); results are presented as mean ± SD (*n*=4); ****: *p* < 0.0001, (c) A representative set of immunofluorescence MIP images after 10-day in vivo oxygenation; i - implantation with oxygenation; ii - implantation without oxygenation; scale bar: 100 μm.

## Conclusion

On site electrocatalytic production of oxygenation demonstrates maintenance of cell viability and therapy production at high cell loadings, enabling more compact cell therapy devices with high dose potential. This is achieved by supplying physiologically required levels of dissolved oxygen in a highly controlled manner (through bioelectronic control of current or voltage), without formation of gas bubbles or generation of hyperoxic (high oxygen) conditions often showing toxic effects. Importantly, this is possible due to the wide operation window for selective oxygen evolution: limiting the side reactions that arise from electrochemistry of species present in the complex cellular environment. In this work, this concept was demonstrated with ARPE-19 at loadings of 60,000 cells/mm^3^ (*ca*. 800 capsules) *in vitro* and *in vivo*, using proper selection of electrocatalyst composition and surface morphology, namely SIROF.

Notably, this work develops the component for oxygenation, which can be readily integrated with other developed technologies for sensing, actuation, communication, and power transfer/generation in an implantable device. For example, while we use batteries in the implant in this work to ease experimentation, the low power requirements (*ca*. 1.25 μW) (especially at low duty cycling) can be readily integrated with wireless power/communications technologies with more favorable safety profiles (i.e., radio frequency/ultrasound/magnetoelectric, etc). Furthermore, ecO_2_ can enable feedback-controlled oxygenation by directly measuring O_2_ concentrations or other metabolic markers to maintain a pre-defined oxygenation setpoint. While this work sets the groundwork for a readily integrated and simplified oxygenation system, future work should address the longevity of the system, exploring non-thin film form factors, especially for application in tissue integration, and expanding the operating winder and selectivity of OER production through the incorporation is novel and existing biomaterials and coatings, including anti-fouling layers, and selective membranes.

This platform for on site, local, and regulated oxygen production suggests broad applicability in biomedical settings. In the cell therapy space, noted herein, ecO_2_ can benefit islet transplantation, as well as cell therapies for onco-therapies, treating metabolic disorders, pain, addiction, and others. Furthermore, given the 14-day period needed to vascularize implanted tissues and cell therapies^47^, temporary oxygenation of tissue transplants can bridge the time from implantation to integration/vascularization from the host recipient. Such an oxygenation platform could be integrated into transport chambers and carriers for long distance transport or donor cells, tissues, or organs. Finally, this device concept can be integrated as a research tool to stress oxygen tension in in vitro organoid models for basic research. As such, a controlled oxygenator that is readily integrated with bioelectronic and biomaterial platforms presents breakthrough opportunities for therapeutics and regenerative engineering applications.

## Acknowledgements

This material is based on research sponsored by 711 Human Performance Wing (HPW) and Defense Advanced Research Projects Agency (DARPA) under agreement number FA8650-21-1-7119. The U.S. Government is authorized to reproduce and distribute reprints for Governmental purposes notwithstanding any copyright notation thereon. The views and conclusions contained herein are those of the authors and should not be interpreted as necessarily representing the official policies or endorsements, either expressed or implied, of 711 Human Performance Wing (HPW) and Defense Advanced Research Projects Agency (DARPA) or the U.S. Government. T.C.-K., S.J., I.L. acknowledge support from Carnegie Mellon University Department of Materials Science and Engineering Materials Characterization Facility supported by Grant MCF-677785. T.C.-K., S.J., I.L. would like to thank Mark Weiler and Dr. Matthew Moneck from the Claire and John Bertucci Nanotechnology Laboratory for assistance with SIROF deposition. J.R., A.S., and X.J., acknowledge Northwestern University’s NUANCE Center, which has received support from the SHyNE Resource (NSF ECCS-2025633), the IIN, and Northwestern’s MRSEC program (NSF DMR-1720139). J.R. and A.S. acknowledge support from Analytical BioNanoTechnology Equipment Core (ANTEC) supported by the Soft and Hybrid Nanotechnology Experimental (SHyNE) Resource (NSF ECCS-2025633) for material characterization and Center for Advanced Microscopy/Nikon Imaging Center (CAM) supported by CCSG P30 CA060553 awarded to the Robert H Lurie Comprehensive Cancer Center for confocal imaging. J.R, T.C-K, A.S., and I.L., acknowledge support from Blackrock Neurotech for SIROF, and wire-bundle bonding for the in-vivo experiments.

## Author Contributions

T.C.-K. and J.R. Conceived the work. I.L., A.S., S.J., G.N., and X.J. performed materials synthesis, characterization, and down selection. I.G, T.S, and I.L. performed finite element modeling of ecO_2_. I.L., A.S., and G.N. and S.J. fabricated and tested electrochemical performance, oxygenation, and side product formation of ecO_2_ devices. I.L., A.S., A.S.M, performed in vitro cell studies and viability studies. S.F., C.F., and O.V. performed cell engineering work and cell encapsulation. A.C., B.R., and J.H. designed and deployed the wireless potentiostat for in vivo experiments. D.J.W., O.E.-R., S.J. and I.L. performed in vivo implantation in rodents. D.S. and A.F. assisted with analysis of cell imaging data and visualization. J.H., D.J.W, O.V., A.F., J.R., and T.C.-K. oversaw the research. I.L., A.S., J.R. and T.C.-K. wrote the manuscript. All authors revised and approved the manuscript.

## Materials and Methods

### ecO_2_ fabrication

Prior to the sputtering of metals, Si/SiO2 (600 nm) wafers were cleaned with sonication in acetone for 10 min, rinsed with IPA and N2 blow dried. The wafers were treated in a barrel etcher with O2 plasma at 100 W for 2 min, immediately followed by spin-coating of LOR10B (MicroChem) at 3,000 rpm for 40 s and baking it at 190 °C for 5 min. After applying photoresist (Shipley, Microposit S1813) at 3,000 rpm for 40 s and baking it at 115 °C for 5 min, UV exposure for photopatterning was carried out with a 4-inch mask in MA6 mask aligner (Karl Suss MA6) for 100 sec. Post-exposure, patterns were developed in 2.6 % tetramethylammonium hydroxide aqueous solution (Shipley, Microposit MF-CD-26 Developer) for 23 sec. The patterned wafers were metalized with a layer of Ti (250 nm) and Pt (450 nm) using DC sputtering at 200 W. Lift-off was conducted in Remover PG (Kayaku Advanced Materials, Inc.) for 30 min at 65 °C. On the metalized wafer, the catalytic array (anode, working electrode) was patterned with Ti (150 nm) and Ir (250 nm) via DC and RF sputtering at 200 W, respectively, while the cathode (counter electrode) remained as intact Pt surface. Reactive sputtering of SIROF was immediately performed after the Ir layer deposition. SIROF was synthesized via reactive sputtering with oxygen plasma^48, 49^. Sputtering was performed in a house-built sputtering system at RF power of 300 W at 10 mTorr with 1:1 mixture of argon and oxygen. A 7 min sputtering resulted in 1.5 μm thick SIROF on an iridium-topped metal stack (Ti 250 nm/Pt 450 nm/Ir 150 nm), which enhances the adhesive properties of SIROF to the metals. Excessive materials were lifted off in Remover PG for 30 min at 65 °C. The patterned arrays were passivated with polymeric window passivation (Kayaku Advanced Materials, Inc, SU-8 3010), followed by hard baking at 180 ℃ for 30 min.

### Electrochemical analysis

All electrochemical analysis was performed in 1 X PBS, unless otherwise noted. Linear sweep voltammetry (LSV) measurement was carried out using a potentiostat (Gamry, Reference 600+) in three-electrode cell setup. To interface the electrode with electrolyte, a 3D-printed reservoir was glued on the catalytic array chip using polydimethylsiloxane (Dow Corning, Sylgard 184). Pt wire and 1 M Ag/AgCl electrodes served as counter and reference electrodes, respectively. LSV curves were collected from 0.7 V to 1.3 V versus 1 M Ag/AgCl at a scan rate of 10 mV s^-1^. Based on the calibration of reference electrodes versus standard hydrogen electrode (SHE), the recorded potential was converted versus reversible hydrogen electrode (RHE) (Equation 1), and overpotential was calculated using Equation 2. Additionally, LSV measurements were also conducted in a two-electrode setup, employing Pt wire as a counter/reference electrode. Onset potential and compliance were determined from the x-intercept of the second derivatives of each LSV curve.

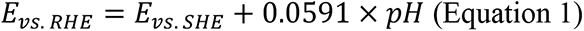

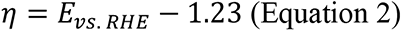

### Oxygen production measurements

An optical oxygen sensor (Presens, PM-PSt7) was used to measure produced oxygen *in vitro*. The sensor needle was replaced with a modified blunt needle, so the sensor was flush with the needle tip. The sensor tip was placed ∼500 µm above the ecO_2_ working electrode using a micromanipulator. The sensor was three-point calibrated by saturating in N_2_, O_2_, and air prior to all measurements. Oxygen sensing experiments were done inside a chamber with a controlled environment under continuous N_2_ purge. Prior to the oxygenation, the electrolyte (∼500 µl) was purged for at least 45 minutes using N_2_ to reach near zero ppm oxygen. Oxygenators were then operated in either chronoamperometry or chronopotentiometry modes for oxygen production. For duty cycle dependance study, a voltage pulse train with varying duty cycle was applied.

### Oxygenation byproducts measurements

For hydrogen peroxide sensing, an ecO_2_ electrode was used in two-electrode configuration in 0.5 ml 1x PBS solution. Prior to the measurement, the electrolyte was purged with N_2_ for ∼1h. The electrode was then subject to oxygen production in chronoamperometry mode for ∼24 hrs. The H_2_O_2_ concentration was measured using a hydrogen peroxide assay kit (Abcam, AB102500) according to the manufacturer’s protocol. Briefly, 48 µL of the assay buffer and standard solutions were added to a 96-well plate, followed by 1 µL each of Oxired Probe and the developer solution. The mixture was agitated for about 5 minutes to ensure thorough mixing. It was then incubated for 1 hour at room temperature before the absorbance was read at 570 nm using a microplate reader (BioTek, Cytation 3). The manufacturer’s standard protocol was used to calibrate the measurements.

For chlorine sensing, commercially available Cl strips (Supelco, 117925) were used. The steps followed are the same as the peroxide sensing. A SIROF electrode with 4 mm^2^ was used as the working electrode, and a platinum wire was used as a counter electrode. After the chronoamperometry measurements, a Cl strip was immersed in the electrolyte for at least 20 s. The color change was then recorded using a camera under consistent lighting conditions. The image was then processed using a custom Matlab program to compare against the standard color values provided by the manufacturer. The greyscale value was used to extrapolate the Cl_2_ concentration. At least four different electrodes were tested for these measurements. All oxygenator measurements were carried out using Keithley 2614 SMU controlled using an open-source software SweepMe. The measured oxygenation data was later corrected using the three-point calibration data obtained earlier.

### On-site cell capsule oxygenation in vitro

To hold media for cell cultivation, PDMS-passivated (Dow chemical, Sylgard 184) 3D-printed media reservoir was applied on catalytic arrays (**Supplementary Figure 8**). Prior to the in vitro experiments, all components that were placed in an incubator were autoclaved (125 ℃ for 30 min) DMEM:F-12 (Gibco, 11320033) formulated with 10 % fetal bovine serum and 1 % pen/strep was employed as a cell culture media. While an in vitro experiment was executed, media was exchanged every 3 days. For additional information see **Supplementary Note 2**.

### Cell viability assay

Cell viability was determined using calcein acetoxymethyl (Calcein AM) and ethidium homodimer (InVitrogen, L3224) for live and dead cells, respectively. Nuclei were labeled using Hoechst 33342 (Thermo Fisher Scientific, 62249). Calcein AM, ethidium homodimer and Hoechst 33342 dyes were added with a final concentration of 2 µM, 4 µM and 1 µM, respectively, to each sample and incubated in a normal incubator for 30 min at 37 °C under 5 % CO_2_. The cells were then washed three times with warm 1 X DPBS. Differential interference contrast (DIC) and scanning laser confocal fluorescence imaging were carried out using an upright confocal microscope and a 20 X/0.50 NA water immersion objective (Nikon). Viability was quantified at least 4 samples per condition and 5 images per sample. Collected images were analyzed by Matlab code to count nuclei, live and dead cells. The viability was calculated as

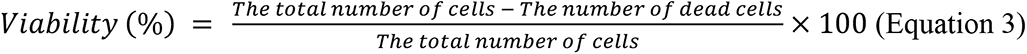

### Leptin cell secretion assay

In order to evaluate peptide production, leptin ELISA kits (Abcam, Human leptin ELISA) were used. Following the manufacturer’s guideline, 3 sets of standard leptin solutions (1,000, 500, 250, 125, 62.5, 31.25,15.63 and 0 pg mL^-1^) were used to generate calibration curves in each round of analysis. The leptin-containing samples were collected 24 hours after complete media exchange to evaluate 24-hour peptide secreting capability. All samples were collected from 4 independent cultivation and technically triplicated with dilution factors of 100X, 500X, 1,000X. The concentration of leptin was measured by recording the absorbance at 450 nm using a plate reader (Molecular Devices, SpectraMax i3X)

### in vivo experiments in rat model

All procedures were approved by the Carnegie Mellon University Institutional Animal Care and Use Committee and the DoD Animal Care and Use Review Office. ecO_2_ was sterilized using ethylene oxide (ETO) sterilization. Maintaining aseptic conditions, ARPE-19 cell capsules were transferred into the cell compartment using a pair of needles. Note that cell capsules were resuspended in fresh media and approximately 800 capsules were loaded. Prior to surgery, a rat was anesthetized using isoflurane in the chamber and it was maintained via nose cone delivery. Shaved rat skin was sanitized using IPA. To minimize mechanical stress on skin and bones, ecO_2_ device was implanted in the abdominal skin pocket. The remote potentiostat was enclosed in 3D-printed housing and placed on the backside to prevent damage from rat’s motion. Those electronics were connected using flexible 16-channel gold wire, which was routed under the skin. For additional information see **Supplementary Note 2**.

