## Supplementary Data for "Electrocatalytic on-site oxygenation for transplanted cell-based-therapies"

### Supplementary Notes

#### Supplementary Note 1. Finite Element Analysis for Oxygen Diffusion Simulation

The COMSOL simulation was constructed in 2-dimensional geometry, which assumed that cross-sectional diffusional profiles were uniform along the axis normal to the plane displayed because of symmetric natures of the designed arrays. The simulations were carried out in a 1 mm high x 2 mm wide rectangle. Oxygen-evolution electrodes were modeled as flat surfaces with pattern spacings according to the electrode designs. The applied physics module, Transport of Diluted Species, limited the oxygen fraction up to their solubility limitation and Fick's Law was selected as a standard diffusion mechanism with the diffusion coefficient of $2x10^{-9}$ $m^2$ $s^{-1}$ for dissolved oxygen in water. The resulting profiles were linearly correlated with applied oxygen flux, the distribution of oxygen was normalized with the oxygen level at the final time stamp.

#### Supplementary Note 2. The Fabrication of $ecO_2$

***In vitro experiment:*** The chip was wire bonded (West Bond, Wedge, Aluminum wire) to an inhouse designed PCB with a 16-pin connector, allowing for individual control of the devices. A custom 3D-printed well was passivated and glued using PDMS (Sylgard 184, Dow chemical) to compartmentalize 8 devices per chip. All components were autoclaved at 125 °C for 30 min prior to the experiment. The alginate encapsulated ARPE-19 cells were dispensed using a pipette to create a uniform monolayer of ~100 capsules in each well. A volume of 250 µl of the basal medium DMEM:F-12 (with 10 % fetal bovine serum and 1 % pen/strep) was then added. The assembled setup was placed in an incubator with an oxygen level of 1% to conduct the experiments. A ribbon connector was used to extend the connection outside the incubator and connect it to potentiostat (Palmsens 4, PalmSens) via 8-channel multiplexer (MUX8-R2, PalmSens). The devices were operated in chronoamperometry mode and voltage was gradually ramped to maintain a constant current. The culture medium was replaced every 72 hours.

***In vivo* experiments in rat model**: A 2×4 array was wire bonded to a PDMS-passivated gold-wire bundle which was then terminated with a 16-pin connector (Omnetics 2207). A PDMS housing with a custom mold was used to enclose the array, with each well holding a 2×4 array. The assembly was then soaked in 70 % ethanol for 24 hours, followed by deionized water (DIW) for another 24 hours, with water exchanges every 8 hours. A commercially available polycarbonate (PCTE) membrane (Sterlitech PCTF 0425100) with 0.4 μm pore size was attached to the top of the well to provide immunoisolation while facilitating nutrient and oxygen exchange. The assembly was then sterilized with EtO and allowed to rest for 48 hours. Prior to the surgical procedure, ARPE-19 cell capsules were transferred into the cell compartment using a pair of needles, maintaining aseptic conditions. The cell capsules were resuspended in fresh media (DMEM:F12 with 10% FBS and 1 % pen/strep), and approximately 800 capsules were loaded.

A rat was anesthetized using isoflurane in a chamber and it was maintained via nose cone delivery. Shaved rat skin was sanitized using IPA. To minimize mechanical stress on skin and bones, $ecO_2$ device was implanted in the abdominal skin pocket. The wire-bundle was then routed under the skin and terminated at the backside to prevent damage from the rat's motion. The Omnetics connector was then connected to the battery-powered remote potentiostat that was enclosed in a 3D-printed housing. Electrochemistry of $ecO_2$ was then wirelessly controlled using a custom-built application on an iOS device.

**Supplementary Note 3. Characterization of SIROF**

***Raman Spectroscopy.*** Raman spectra of SIROF were collected using NT-MDT Spectra with 532-nm excitation through 100x/0.7 NA objective. The spectra were recorded with a 0.5 neutral density (ND) filter and 30-second acquisition time, acquired from 10 random locations of 4 independent samples. While detected $E_g$ (*ca*. 550 cm$^{-1}$), $B_{2g}$ and $A_{1g}$ (*ca*. 720 cm$^{-1}$) modes indicate crystalline synthesized SIROF, overwrapped $B_{2g}$ and $A_{1g}$ peaks showed amorphous domains also existed in the materials.[3-5] In addition to main peaks, weak mode at ~365 cm$^{-1}$ also supported the presence of amorphous iridium oxide, derived from stretches of disordered iridium oxide structure.[4]

***Grazing Incidence X-ray diffractometry (GIXRD).*** To investigate crystallographic features of SIROF, we acquired the XRD spectra using grazing incidence to avoid interference from materials stacked under the films. The GIXRD patterns were collected using Malvern Panalytical Empyrean XRD with a wavelength of 0.154 nm (Cu Kα) and a grazing angle of 3° at 45 kV and 40 mA. The scan step size was 0.02° with 1 sec per scan. The diffraction patterns at 28.314°, 34.716° and 40.350° are corresponding to (110), (101) and (200) plane of rutile $IrO_2$ lattice, respectively, which implied that dendritic SIROF nanostructures were composed of nanocrystals.[5-7]

***X-ray photoemission spectroscopy (XPS).*** SIROF surface chemical composition was investigated using XPS. The spectra were acquired using Thermo Scientific ESCALAB 250Xi equipped with a monochromatic KR Al X-ray source (spot size: 500 μm). Survey spectra were collected with a 120-eV pass energy, 15-ms dwell time and 1-eV step size. High-resolution scans were performed in Ir 4f, O 1s and C 1s regions with 0.1-eV step size, and C-C peak was utilized for calibration as 284.8 eV. The trace level of Si signals implied that collected XPS spectra were free from the substrate (Si/$SiO_2$ (600 nm thermal oxide) wafer). The recorded XPS signals were analyzed using Avantage (Thermo Scientific) software. Ir 4f doublet peaks were deconvoluted using asymmetric curve fitting with Ir (IV) and its first satellite signal.[8] Corresponding to electrochemical analysis displaying the absence of the transition from Ir (III) to Ir (IV), iridium species in SIROF represented +4 oxidation state, which is active for the electrocatalysis of oxygen evolution reaction.[9] Note that including Ir(III) species did not create reasonable results, which indicated +4 oxidation state is the only chemical status that existed in SIROF. A high-resolution scan of O 1s regime indicated the synthesized SIROF catalysts implied that synthesized SIROF possessed oxygen vacancies as well was lattice oxygen.[10,11] Because while oxygen vacancy signals are originated from amorphous iridium oxide ($IrO_X$), lattice oxygen is detected in $IrO_2$ crystal, the chemical status of oxygen in SIROF provides a hint of semi-crystalline characteristics. Combined with the structural analyses showing both amorphous characteristics (Raman spectra) and crystalline (GIXRD), the mixed signals from lattice and vacancy oxygen in XPS analysis substantiated that SIROF has semi-crystalline natures. As reported that oxygen vacancies are key components in oxygen evolution mechanism of iridium oxide catalysts,[12] the existence of Ir (IV) states and oxygen vacancies indicated that synthesized SIROF has catalytic activity in OER.

***Scanning Electron Microscopy (SEM).*** SEM images were acquired by using a field emission gun SEM (Quanta 600, FEI). All images were acquired with high-resolution (2048 x 1768 pixels) at an accelerating voltage of 1 kV with a working distance of 5 mm. No additional conductive coating was applied to any of the samples for the imaging.

**Supplementary Note 4. Engineered ARPE-19 Alginate Cell Capsules**

***Cell Engineering.*** To engineer cells expressing leptin APRE-19, cells were transfected with a CAG-Leptin plasmid using a piggybac transposase helper plasmid. ARPE-19 cells were seeded at a density of 300k cells/well of a 6 well plate and incubated overnight at 37℃ in a 5% CO2 humidified atmosphere. Cells were then transfected using Lipoectamine 3000 according to the manufacturer's protocol (Catalog no. L3000015 Thermo Fisher Scientific) with the leptin and helper plasmid in a ratio of 2:1. Beginning 24 hours after transfection the cells were selected for transgene expression using media with 2 μg/mL of puromycin (Catalog no. A1113803 Thermo Fisher Scientific).

***ARPE-19 alginate cell capsule fabrication.*** To prepare for alginate encapsulation alginate was dissolved at 1.4% (w/v) in 0.8% saline and sterile-filtered through a 0.2-μm syringe filter and all buffers were prepared and sterilized by autoclaving and sterile filtering through a 0.2-μm vacuum filter. Cells were trypsinized and collected before being washed three times with calcium-free Krebs solution (4.7 mM KCl, 25 mM HEPES, 1.2 mM KH2PO4, 1.2 mM MgSO4·7H2O, 135 mM NaCl) by centrifuging the cells at 250 g for 5 minutes.

After the third wash, the cells were resuspended in alginate at a concentration of 60k cells mm-3. Encapsulation was done via electrospraying custom-built electrostatic spraying device. The device consisted of a syringe pumps (Harvard Apparatus), a 30 gauge blunt tip needle and a voltage generator (Gamma High Voltage) that was attached to the tip of the needle and grounded to a glass dish containing a cross-linking bath (20 mM barium, 5% mannitol). Cell ladlen alginate droplets were expelled from the needle into the cross-linking solution at a rate of 3 ml/hour. The size of the capsules was maintained by adjusting the voltage on the generator with a voltage of ~6.14 kV applied to produce capsules that were 300 μm in diameter. Capsules were incubated in the cross-linking bath for 10 min to ensure complete cross-linking. They were subsequently washed three times with Hepes buffer (0.132 M NaCl, 4.7mM KCl, 1.2mM $MgCl2 \cdot 6H2O$, 25 mM HEPES) three times and transferred to a flask containing complete cell culture medium (Dulbecco's modified Eagle's medium (DMEM/F-12), with 10% fetal bovine serum (FBS) and 1% antibiotic-antimycotic (AA)) and maintained with standard cell culture techniques prior to experimental setup.

**Supplementary Note 5. Remote Potentiostat**

*Potentiometry circuit equipped for wireless communication.* A compact (12 mm x 20 mm) printed circuit board was developed to provide the necessary voltage to the oxygenation electrode and to enable real-time in vivo monitoring of oxygen generation. The circuit features a Bluetooth Low Energy (BLE) enabled microcontroller, chosen for its low power consumption, and a dual-channel 12-bit buffered digital-to-analog converter (DAC) that allows for precise biasing of the catalysts. Four oxygen-generating channels and two calibration channels are connected in series to the DAC outputs through digitally-controlled single-pole double-throw (SPDT) analog switches. The circuit is powered by a lithium polymer battery and supporting circuitry. One DAC channel continuously supplies the catalysts with a known voltage (the "oxygenation output"), while the other is wired in a force-sense configuration with a feedback resistor (the "measurement output"). The microcontroller selects the DAC output for each of the four oxygen-generating channels.

During normal operation, all channels are connected to the oxygenation output. Periodically, the microcontroller switches each channel in sequence to the measurement output, programmed to produce the same voltage as the oxygenation output. After a delay, the microcontroller's analog-to-digital converter (ADC) reads the voltage across the feedback resistor and calculates the current flowing into each channel. The calibration channels connect to precision (0.5%) resistors, facilitating measurement calibration throughout the experiment as conditions change. The voltage, measurement frequency, and delay can be wirelessly programmed in real-time using a companion mobile application. The microcontroller transfers measurement data to the application once its buffer is full, allowing the operator to view and adjust parameters for continued operation.

### Supplementary Tables

**Supplementary Table 1. Data summary of XPS elemental analysis.** Samples were prepared on $Si/SiO_2$ (600 nm) wafer. Given trace level of Si signal, collected atomic fractions were free from the substrates. Results are presented as mean ± SD (*n*=4).

| | Element | Atomic ratio (%) |
|---|---|---|
| **Sample #1** | Ir | 23.6 |
| | O | 64.3 |
| | C | 12.1 |
| | Si | trace (< 0.5) |
| **Sample #2** | Ir | 22.5 |
| | O | 63.9 |
| | C | 13.6 |
| | Si | trace (< 0.5) |
| **Sample #3** | Ir | 23.4 |
| | O | 65.8 |
| | C | 10.9 |
| | Si | trace (< 0.5) |
| **Sample #4** | Ir | 21.7 |
| | O | 62.9 |
| | C | 15.4 |
| | Si | trace (< 0.5) |
| **Average** | Ir | 22.8 ± 0.76 |
| | O | 64.23 ± 1.04 |
| | C | 13.0 ± 1.69 |
| | Si | trace (< 0.5) |

**Supplementary Table 2. Data summary for XPS of SIROF in C 1s region.** C-C state at 284.4 eV in C 1s region was utilized for chemical shift calibration. Results are presented as mean ± SD (*n*=4).

| **Peak** | **Sample** | **State** | **Peak BE** (eV) | **FWHM** (eV) | **Ratio** |
|---|---|---|---|---|---|
| **C 1s** | **Sample #1** | C-C | 284.8 | 1.7 | 100 |
| | **Sample #2** | C-C | 284.8 | 1.8 | 100 |
| | **Sample #3** | C-C | 284.8 | 1.8 | 100 |
| | **Sample #4** | C-C | 284.8 | 1.7 | 100 |
| | **Average** | C-C | 284.8 ± 0 | 1.8 ± 0.05 | 100 ± 0 |

**Supplementary Table 3. Data summary for XPS of SIROF in Ir 4f region.** Results are presented as mean ± SD (*n*=4).

| **Peak** | **Sample** | **State** | **Peak BE** (eV) | **FWHM** (eV) | **Ratio** |
|---|---|---|---|---|---|
| **Ir 4f**[8] | **Sample #1** | Ir(IV) 7/2 | 62.2 | 1.8 | 39.6 |
| | | Ir(IV) 5/2 | 65.2 | 1.8 | 39.8 |
| | | Ir (IV) 7/2 sat. | 63.4 | 2.9 | 10.3 |
| | | Ir (IV) 5/2 sat. | 66.6 | 2.9 | 10.4 |
| | **Sample #2** | Ir(IV) 7/2 | 62.2 | 1.8 | 38.5 |
| | | Ir(IV) 5/2 | 65.2 | 1.8 | 38.7 |
| | | Ir (IV) 7/2 sat. | 63.4 | 3.0 | 11.4 |
| | | Ir (IV) 5/2 sat. | 66.6 | 3.0 | 11.4 |
| | **Sample #3** | Ir(IV) 7/2 | 62.3 | 1.8 | 38.3 |
| | | Ir(IV) 5/2 | 65.3 | 1.8 | 38.3 |
| | | Ir (IV) 7/2 sat. | 63.4 | 2.9 | 11.7 |
| | | Ir (IV) 5/2 sat. | 66.6 | 2.9 | 11.7 |
| | **Sample #4** | Ir(IV) 7/2 | 62.1 | 1.8 | 38.7 |
| | | Ir(IV) 5/2 | 65.1 | 1.8 | 38.7 |
| | | Ir (IV) 7/2 sat. | 63.2 | 3.0 | 11.3 |
| | | Ir (IV) 5/2 sat. | 66.4 | 3.0 | 11.3 |
| | **Average** | Ir(IV) 7/2 | 62.2 ± 0.07 | 1.8 ± 0 | 38.8 ± 0.50 |
| | | Ir(IV) 5/2 | 65.2 ± 0.07 | 1.8 ± 0 | 38.8 ± 0.56 |
| | | Ir (IV) 7/2 sat. | 63.4 ± 0.09 | 3.0 ± 0.05 | 11.2 ± 0.53 |
| | | Ir (IV) 5/2 sat. | 66.6 ±0.09 | 3.0 ± 0.05 | 11.2 ± 0.49 |

**Supplementary Table 4. Data summary for XPS of SIROF in O 1s region.** Results are presented as mean ± SD (*n*=4).

| **Peak** | **Sample** | **State** | **Peak BE** (eV) | **FWHM** (eV) | **Ratio** |
|---|---|---|---|---|---|
| **O 1s**[10,11] | **Sample #1** | $IrO_2$ (lattice) | 530.5 | 1.5 | 40.8 |
| | | $IrO_X$ (vacancy) | 531.6 | 1.5 | 29.7 |
| | | IrOH | 532.6 | 1.5 | 18.5 |
| | | C-O | 533.7 | 1.5 | 8.4 |
| | | $H_2O$, abs. | 535.1 | 1.5 | 2.7 |
| | **Sample #2** | $IrO_2$ (lattice) | 530.6 | 1.4 | 40.3 |
| | | $IrO_X$ (vacancy) | 531.7 | 1.4 | 30.7 |
| | | IrOH | 532.7 | 1.4 | 18.7 |
| | | C-O | 533.7 | 1.4 | 7.9 |
| | | $H_2O$, abs. | 535.0 | 1.4 | 2.3 |
| | **Sample #3** | $IrO_2$ (lattice) | 530.5 | 1.5 | 38.2 |
| | | $IrO_X$ (vacancy) | 531.6 | 1.5 | 27.4 |
| | | IrOH | 532.5 | 1.5 | 15.0 |
| | | C-O | 533.5 | 1.5 | 8.8 |
| | | $H_2O$, abs. | 534.9 | 1.5 | 3.0 |
| | **Sample #4** | $IrO_2$ (lattice) | 530.5 | 1.5 | 41.7 |
| | | $IrO_X$ (vacancy) | 531.6 | 1.5 | 32.1 |
| | | IrOH | 532.6 | 1.5 | 16.0 |
| | | C-O | 533.8 | 1.5 | 7.8 |
| | | $H_2O$, abs. | 535.3 | 1.5 | 2.4 |
| | **Average** | $IrO_2$ (lattice) | 530.5 ± 0.04 | 1.5 ± 0.04 | 40.3 ± 1.29 |
| | | $IrO_X$ (vacancy) | 531.6 ± 0.04 | 1.5 ± 0.04 | 30.0 ± 1.71 |
| | | IrOH | 532.6 ± 0.07 | 1.5 ± 0.04 | 17.1 ± 1.59 |
| | | C-O | 533.7 ± 0.11 | 1.5 ± 0.04 | 8.2 ± 0.40 |
| | | $H_2O$, abs. | 535.1 ± 0.15 | 1.5 ± 0.04 | 2.6 ± 0.27 |

### Supplementary Figures

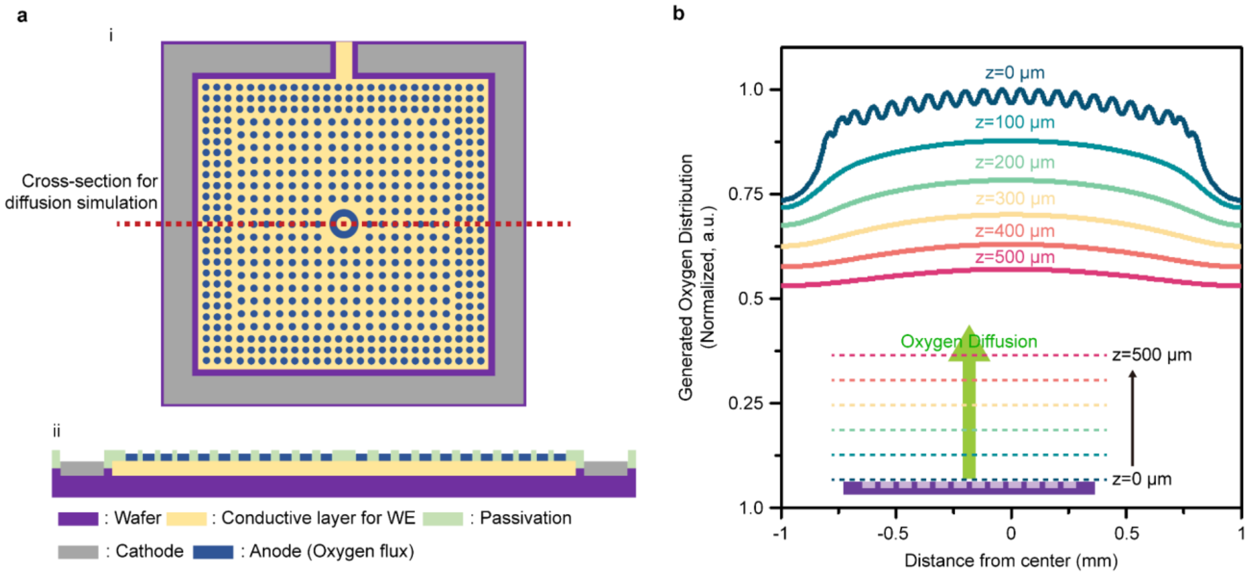


**Supplementary Figure 1. Simulated cross-sectional oxygen profiles**. (a) The designed electrode for diffusion simulation; i – top view of the electrode; ii – cross-sectional view of the electrode at the center of the electrode. (b) The simulated oxygen profiles at different z-positions. The oxygen levels were normalized with the value at z=0 μm; inset: a schematic illustration describing how the profiles were exported and plotted.

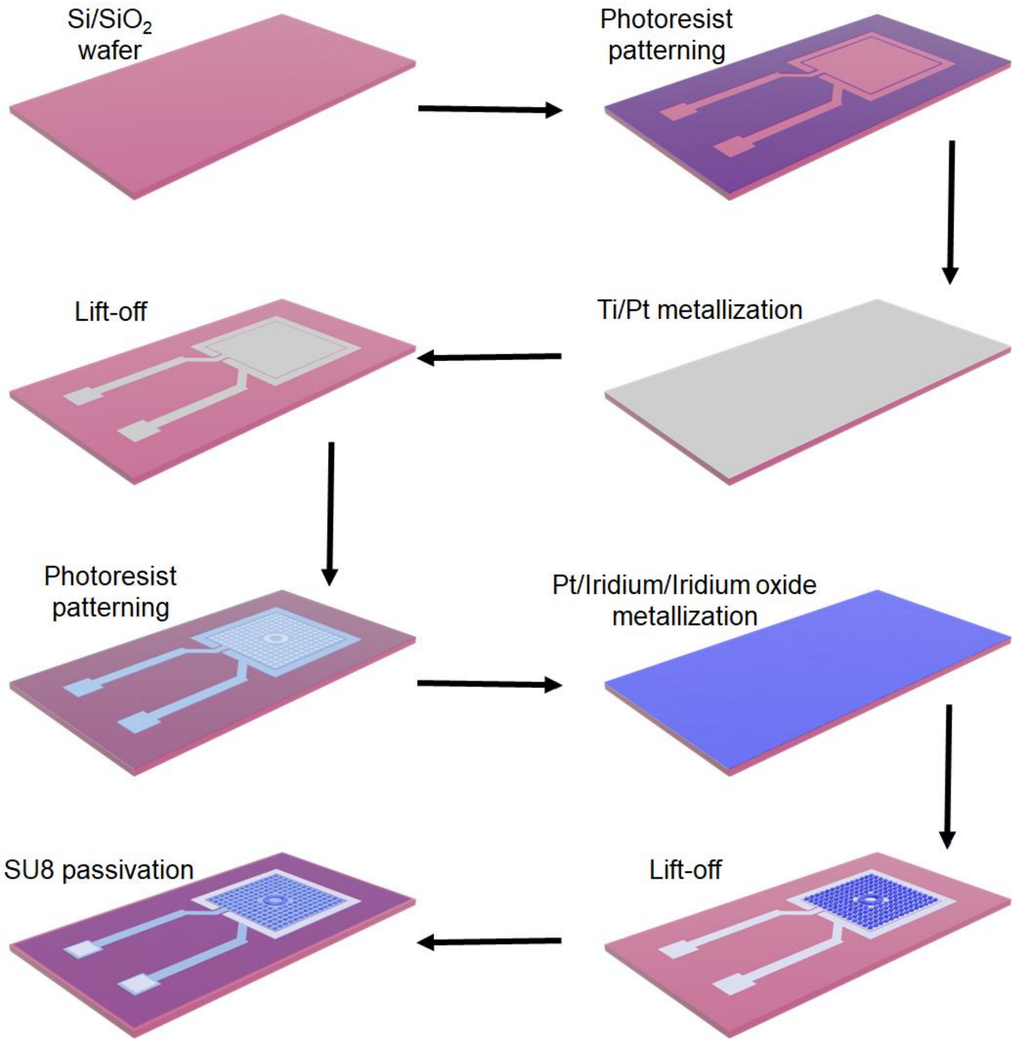


**Supplementary Figure 2. ec$O_2$ microelectrode fabrication scheme.** ec$O_2$ microelectrodes were fabricated on a silicon wafer with 600 nm thick thermally grown $SiO_2$. Detailed description is provided in materials and methods.

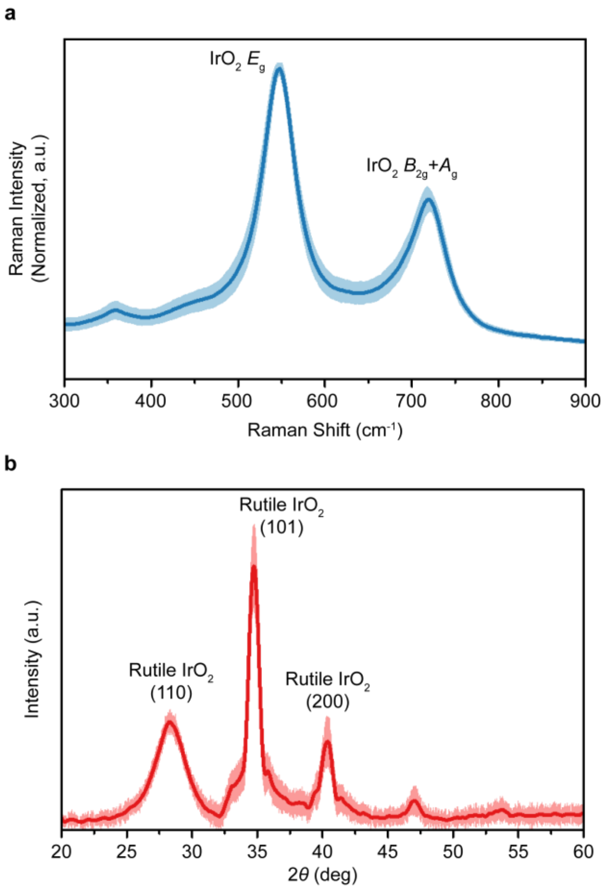


**Supplementary Figure 3. Structural analysis of the SIROF catalyst**. (a) Raman spectroscopy (blue) and (b) Grazing incidence X-ray diffractometry (GIXRD) (red). Results are presented as mean ± SD ($n$=4-6).

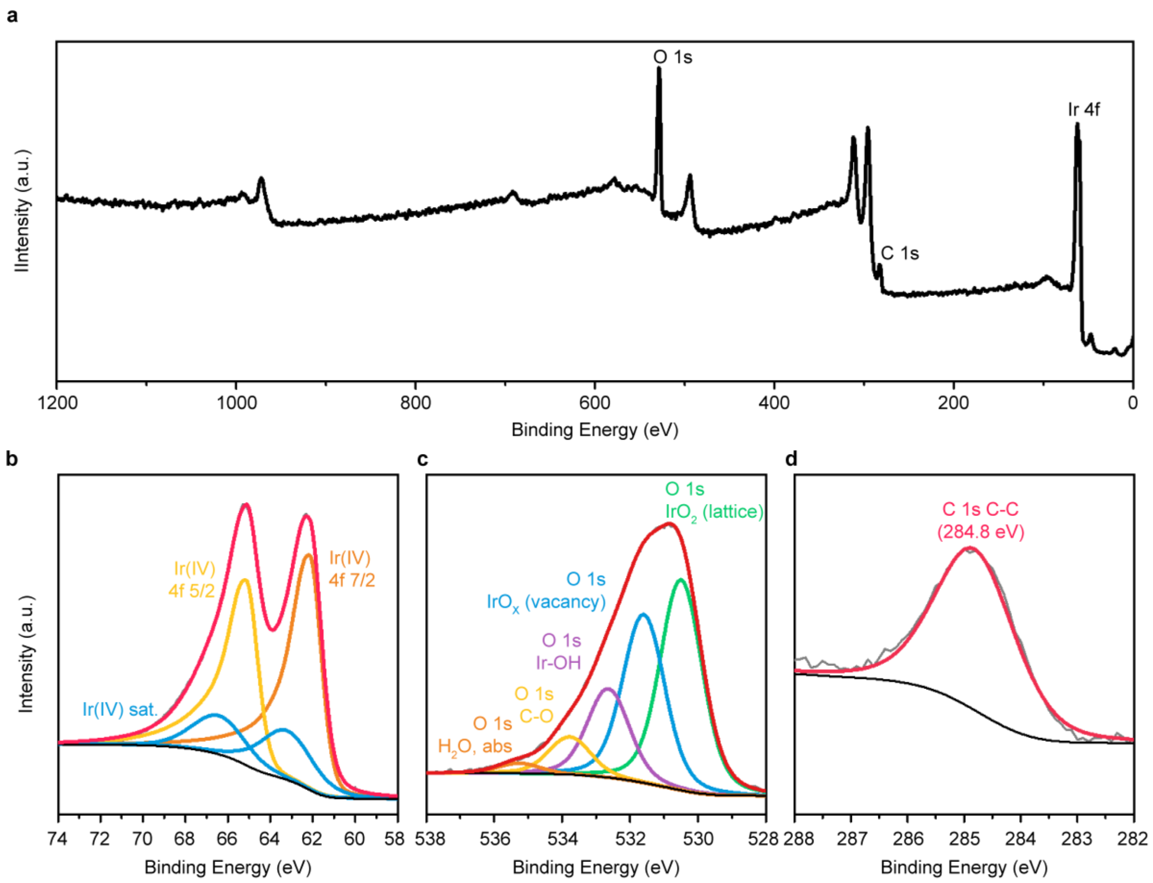


**Supplementary Figure 4. X-ray photoemission spectroscopy (XPS) analysis of SIROF**. Representative XPS deconvolution. (a) survey scan; step size – 1 eV (b) Ir 4f; asymmetric doublet curve fitting for Ir(IV) was applied (c) O 1s; (d) C 1s; C-C peak at 284.8 eV was utilized for chemical shift calibration (*n*=4); step size for high-resolution scan – 0.1 eV.

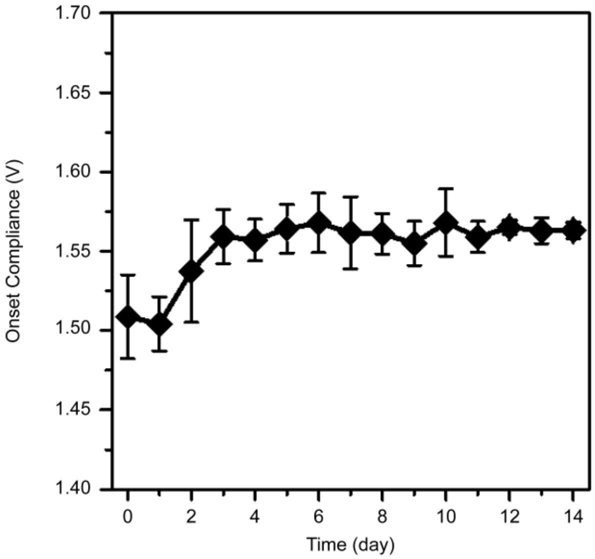


**Supplementary Figure 5. ec$O_2$ onset compliance.** Onset values were calculated from 2 electrodes LSV curves which were collected by every 24 hours over 14 days during chronoamperometry at 1.7 V in 1X PBS. Results are represented as mean ± SD (*n*=5)

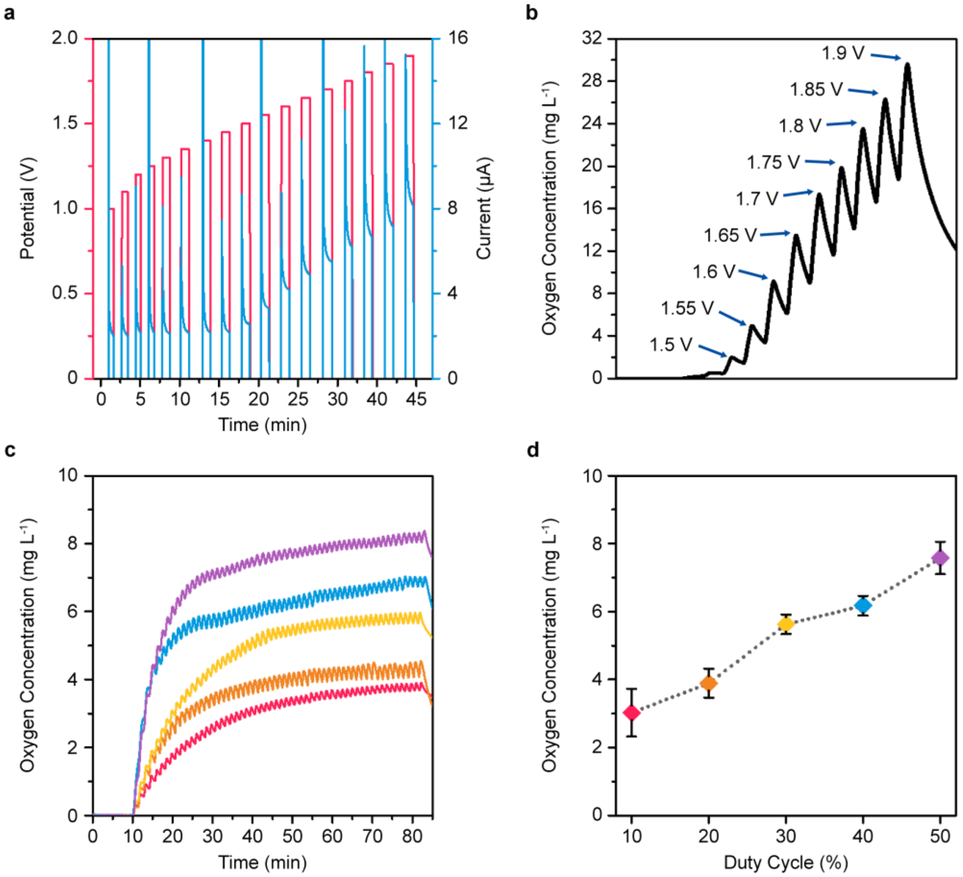


**Supplementary Figure 6. Potentiometric electrochemical oxygen generation.** (a) A representative recorded current profile (blue) from applied potential (red). (b) A representative measured oxygen concentration profiles with 30-sec potential pulses. (c) A representative produced oxygen profiles by various duty cycles; 10 % (red), 20 % (yellow), 30 % (green), 40 % (blue) and 50 % (purple) at the compliance of 1.7 V. (d) measured oxygen concentration after 60 min oxygen production with varied duty cycles. Results are presented as mean ± SD (*n*=3).

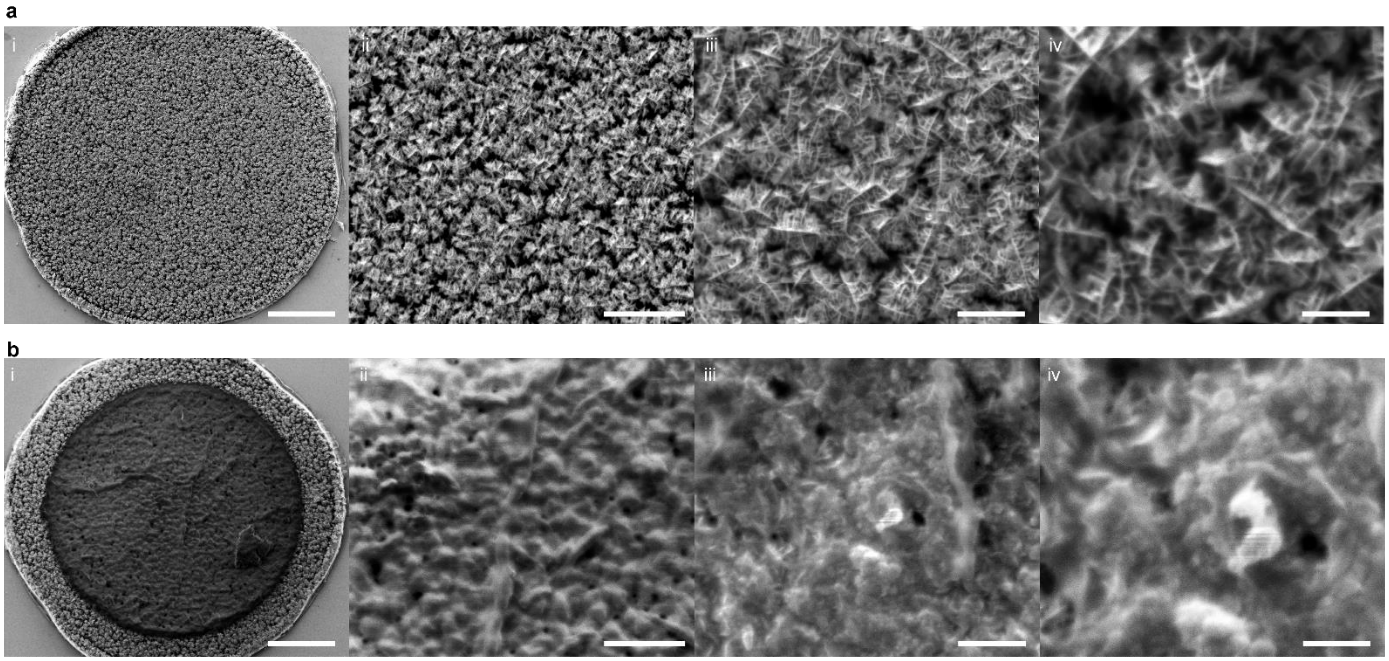


**Supplementary Figure 7. ecO2 before and after 21-day oxygen evolution reaction**. Representative SEM image (a) before and (b) after electrochemical oxygen evolution for 21 days with 100 % duty cycle load; scale bars: i – 10 µm; ii – 2 µm; iii – 1 µm; iv – 500 nm. All images were collected at an accelerating voltage of 1 kV with a working distance of 5 mm.

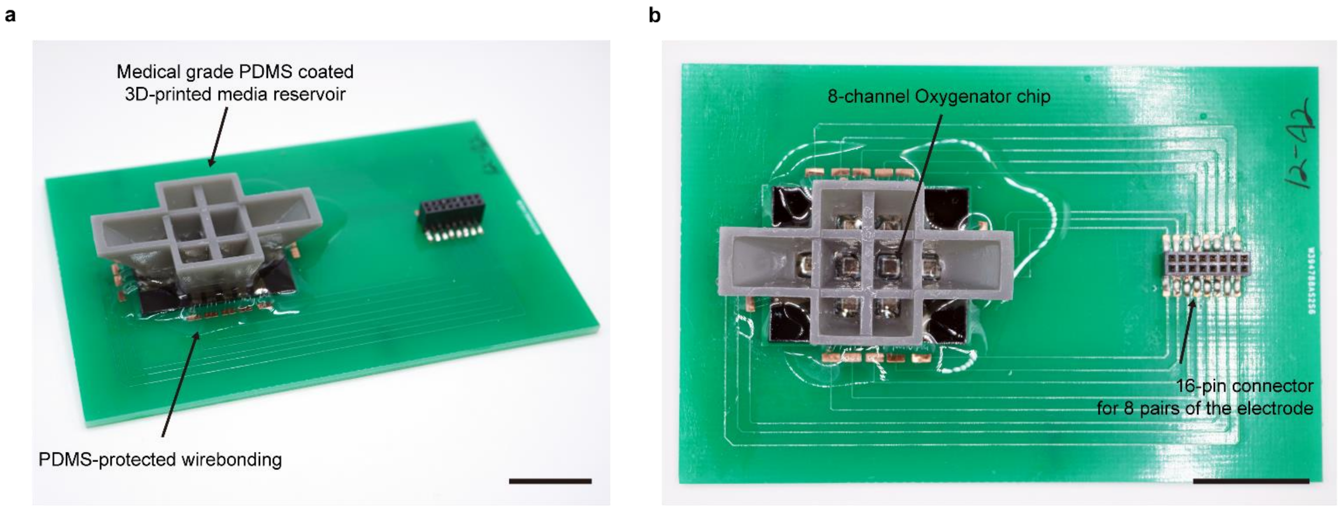


**Supplementary Figure 8. $ecO_2$ *in vitro* device.** Representative images of (a) side view and (b) top view of the in vitro $ecO_2$. Scale bars are 2 cm.

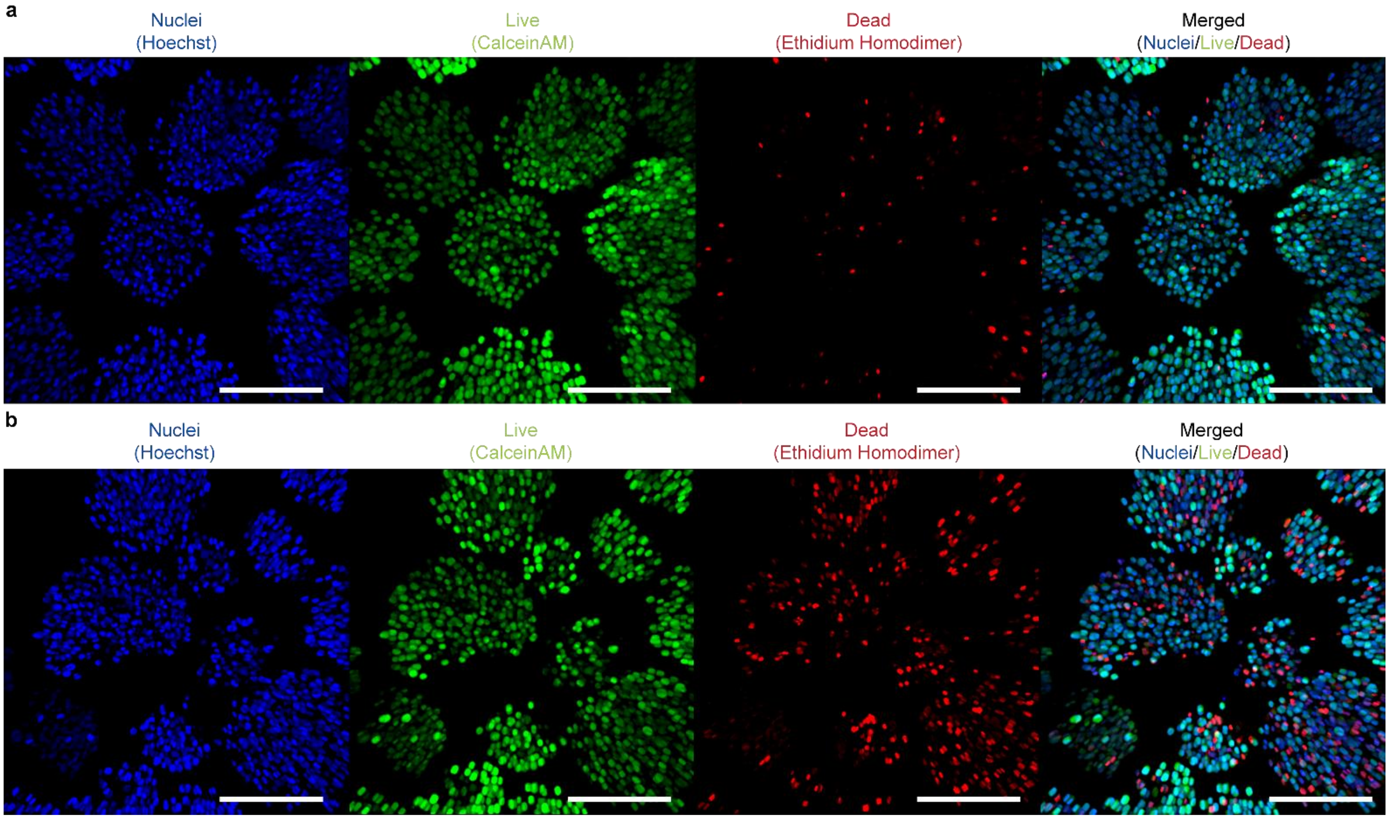


**Supplementary Figure 9. Live/dead assay fluorescence images from 3-day *in vitro***. Representative image for (a) with and (b) without oxygenation. All scale bars are corresponding to 200 μm. Green: CalceinAM (Live), Red: Ethidium homodimer (Dead), Blue: Hoechst (Nuclei).

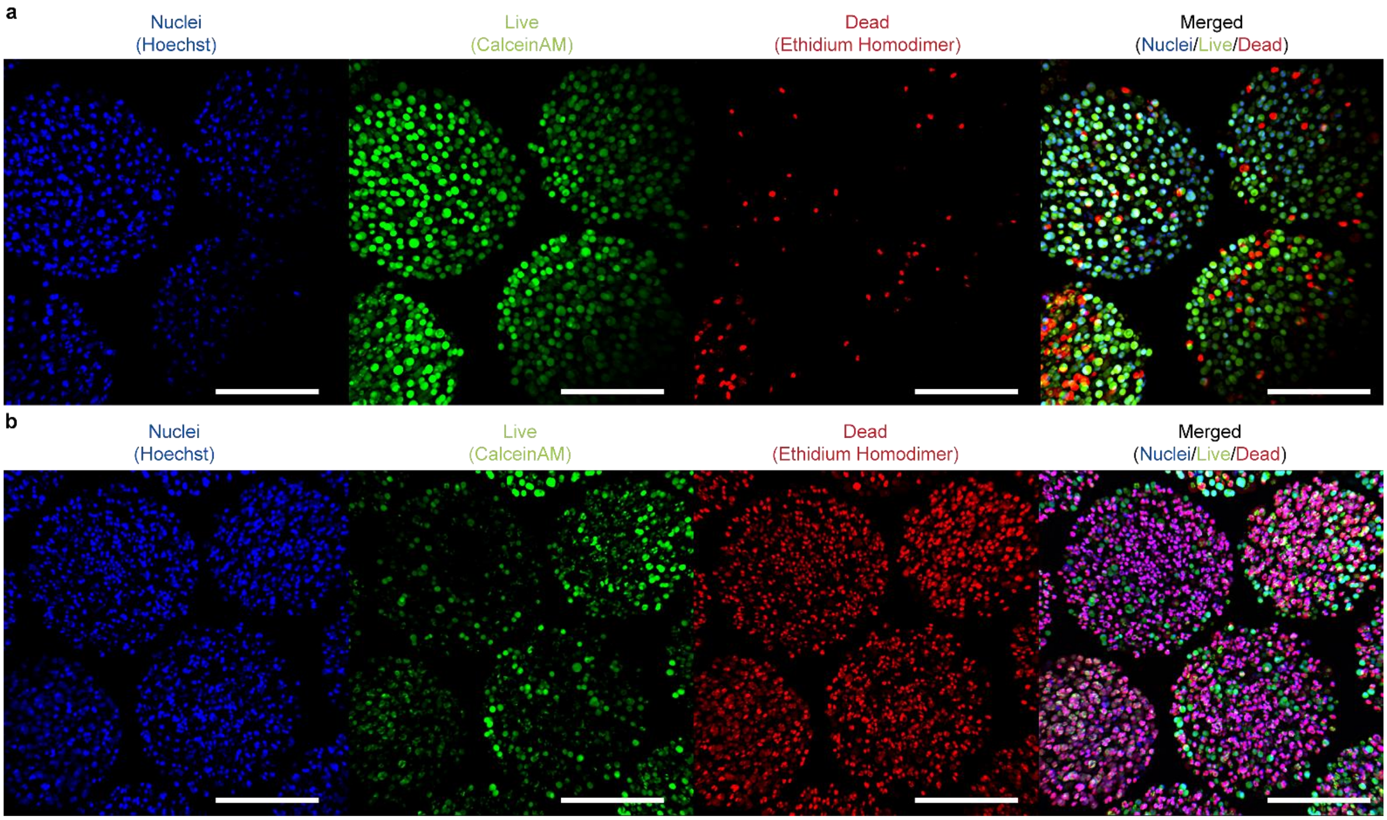


**Supplementary Figure 10. Live/dead assay fluorescence images from 10-day *in vitro***. Representative image for (a) with and (b) without oxygenation. All scale bars are corresponding to 200 μm. Green: CalceinAM (Live), Red: Ethidium homodimer (Dead), Blue: Hoechst (Nuclei).

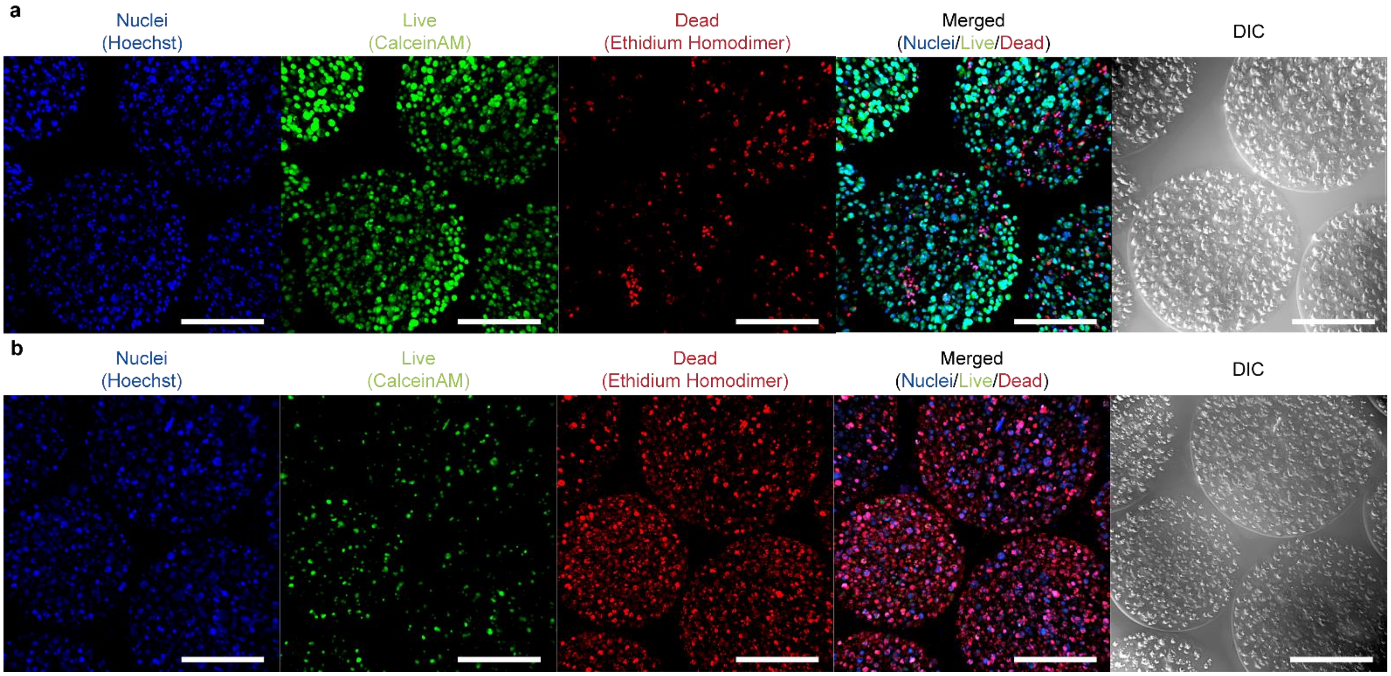


**Supplementary Figure 11. Live/dead assay fluorescence images from 21-day *in vitro***. Representative image for (a) with and (b) without oxygenation. All scale bars are corresponding to 200 μm. Green: CalceinAM (Live), Red: Ethidium homodimer (Dead), Blue: Hoechst (Nuclei), DIC: differential interference contrast images (Cell capsule).

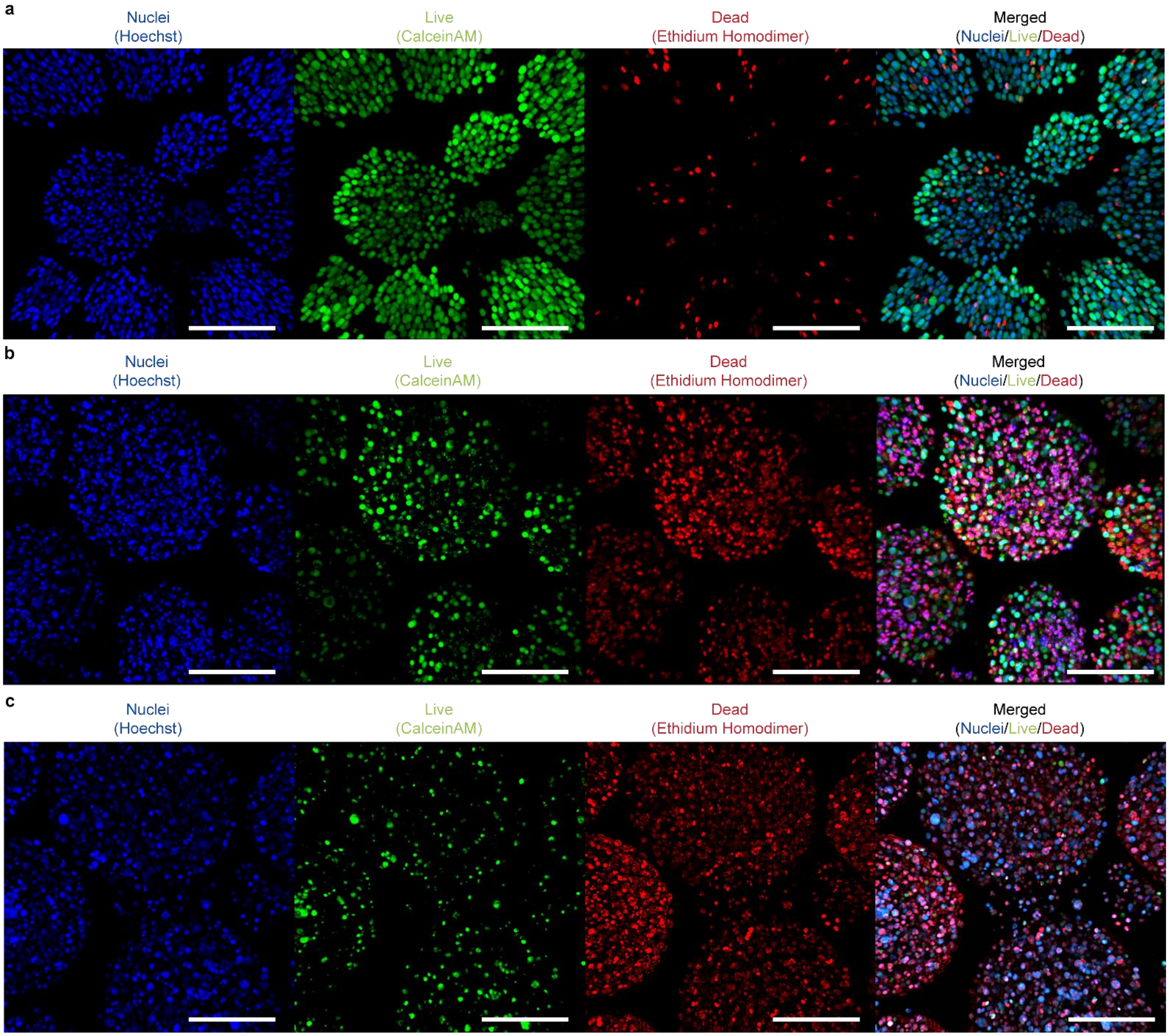


**Supplementary Figure 12. Live/dead assay fluorescence images from normoxic incubation (20% $O_2$) without oxygenation.** Representative images for (a) 3, (b) 10 and (c) 21 days. All scale bars are corresponding to 200 μm. Green: CalceinAM (Live), Red: Ethidium homodimer (Dead), Blue: Hoechst (Nuclei).

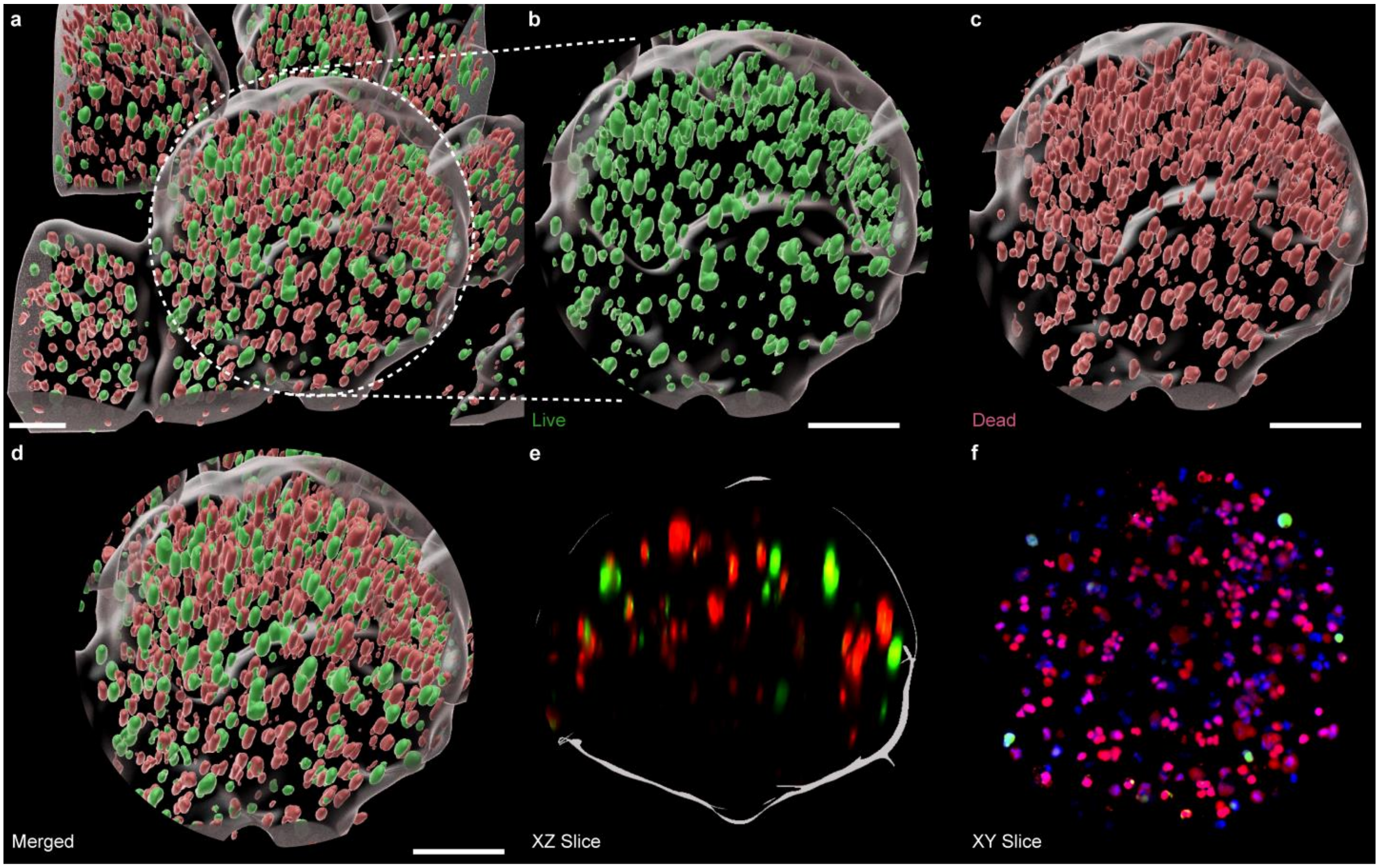


**Supplementary Figure 13. 3D-rendered fluorescence images of cell capsules after 21-day normoxic incubation**. (a) 3D-reconstructed z-stack live/dead fluorescence images, (b) Live and (c) Dead cells in the white dashes circle marked capsule. A cross-section view in (d) XZ plane and in (e) XY plane. Scale bars are 100 µm. Green: CalceinAM (Live), Red: Ethidium homodimer (Dead), Blue: Hoechst (Nuclei).

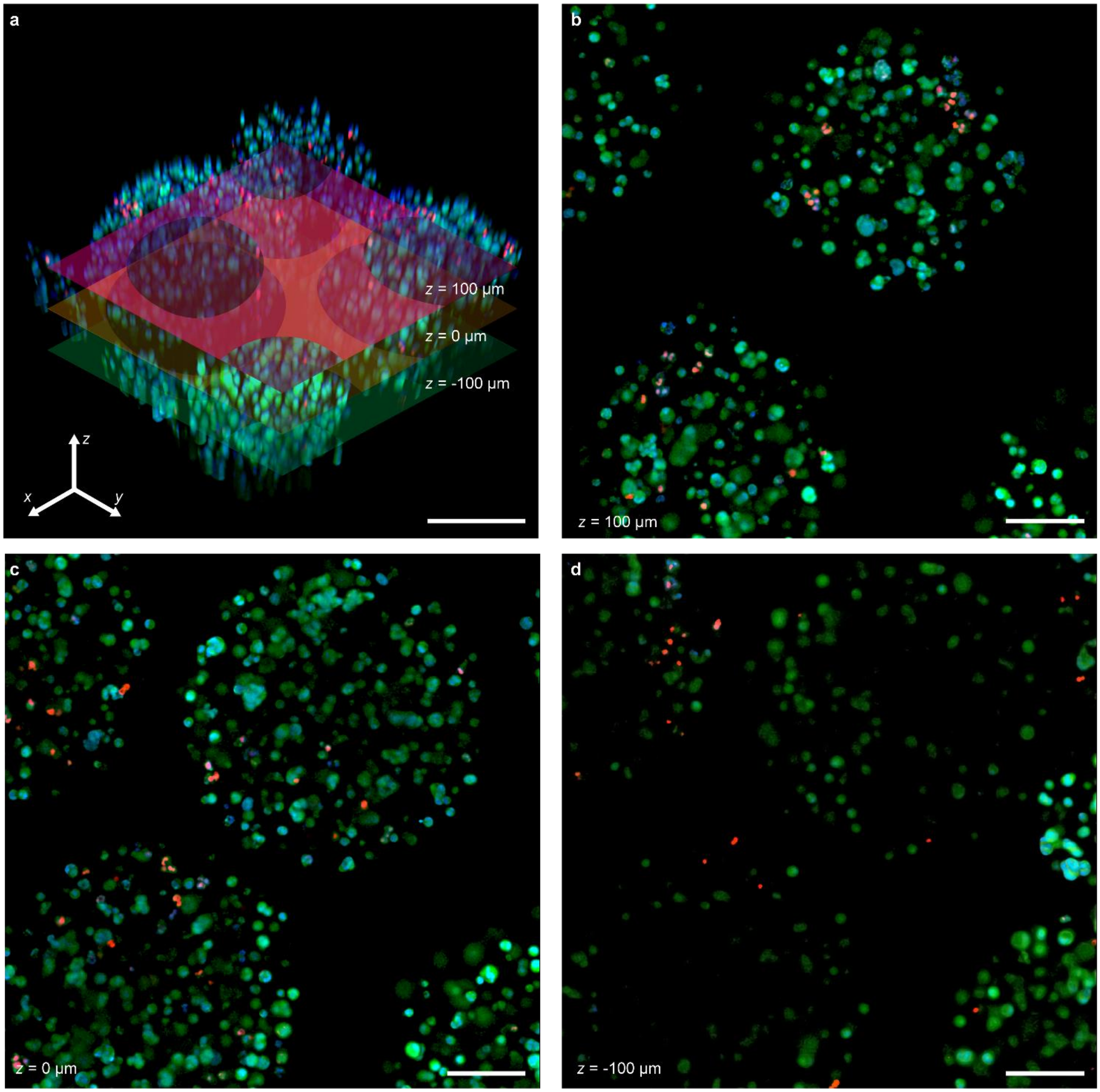


**Supplementary Figure 14. Z-stack analysis of 21-day oxygenation**. (a) 3D-reconstructed z-stacked images and the location of each presented image. (b) z=100 µm; (c) z=0 µm; (d) z=-100 µm. Scale bars are 100 µm. Green: CalceinAM (Live), Red: Ethidium homodimer (Dead), Blue: Hoechst (Nuclei).

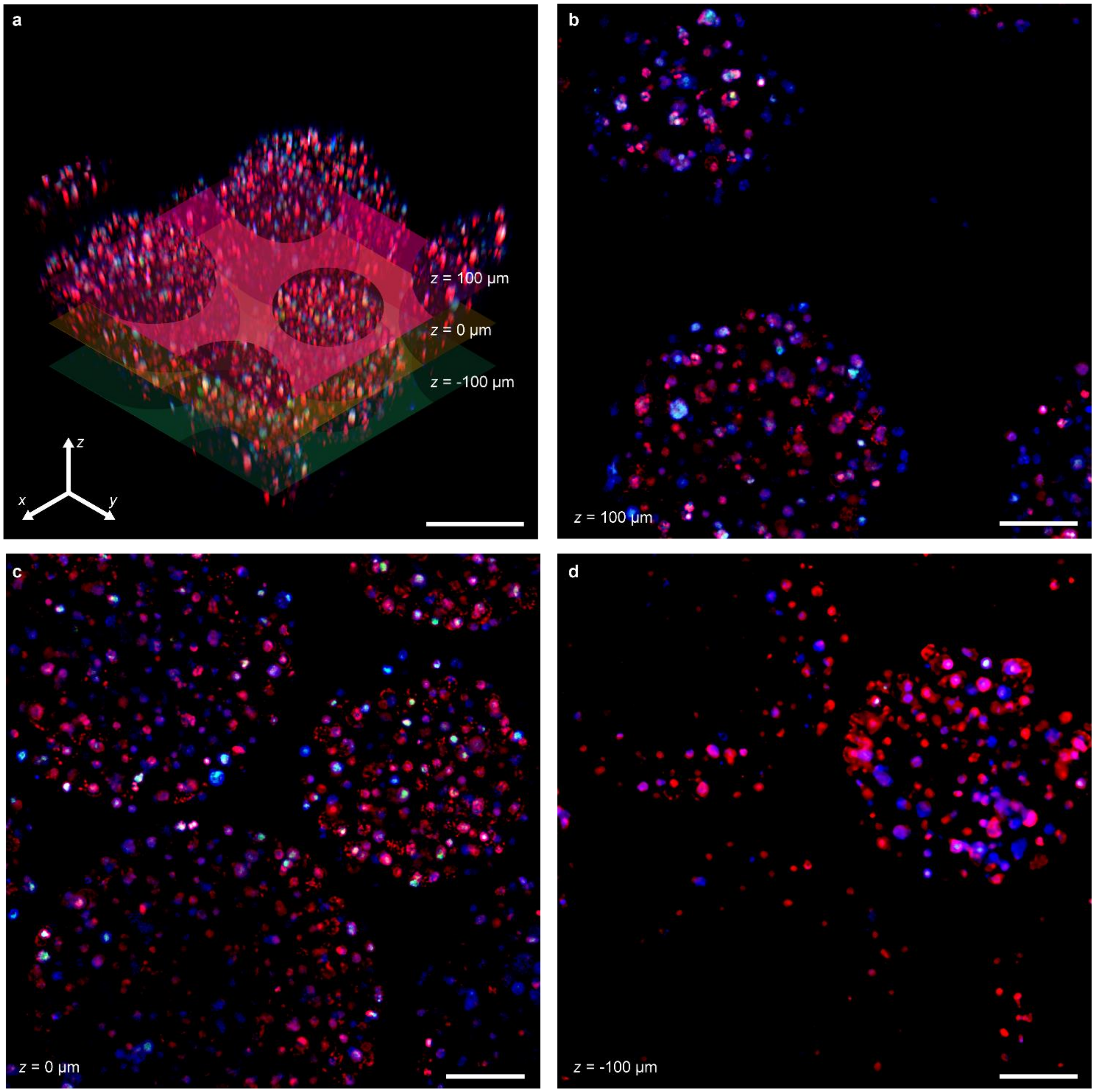


**Supplementary Figure 15. Z-stack analysis of 21-day hypoxia (1% $O_2$) control**; (a) 3D-reconstructed z-stacked images and the location of each presented image. (b) z=100 μm; (c) z=0 μm; (d) z=-100 μm; Scale bars are 100 μm. Green: CalceinAM (Live), Red: Ethidium homodimer (Dead), Blue: Hoechst (Nuclei).

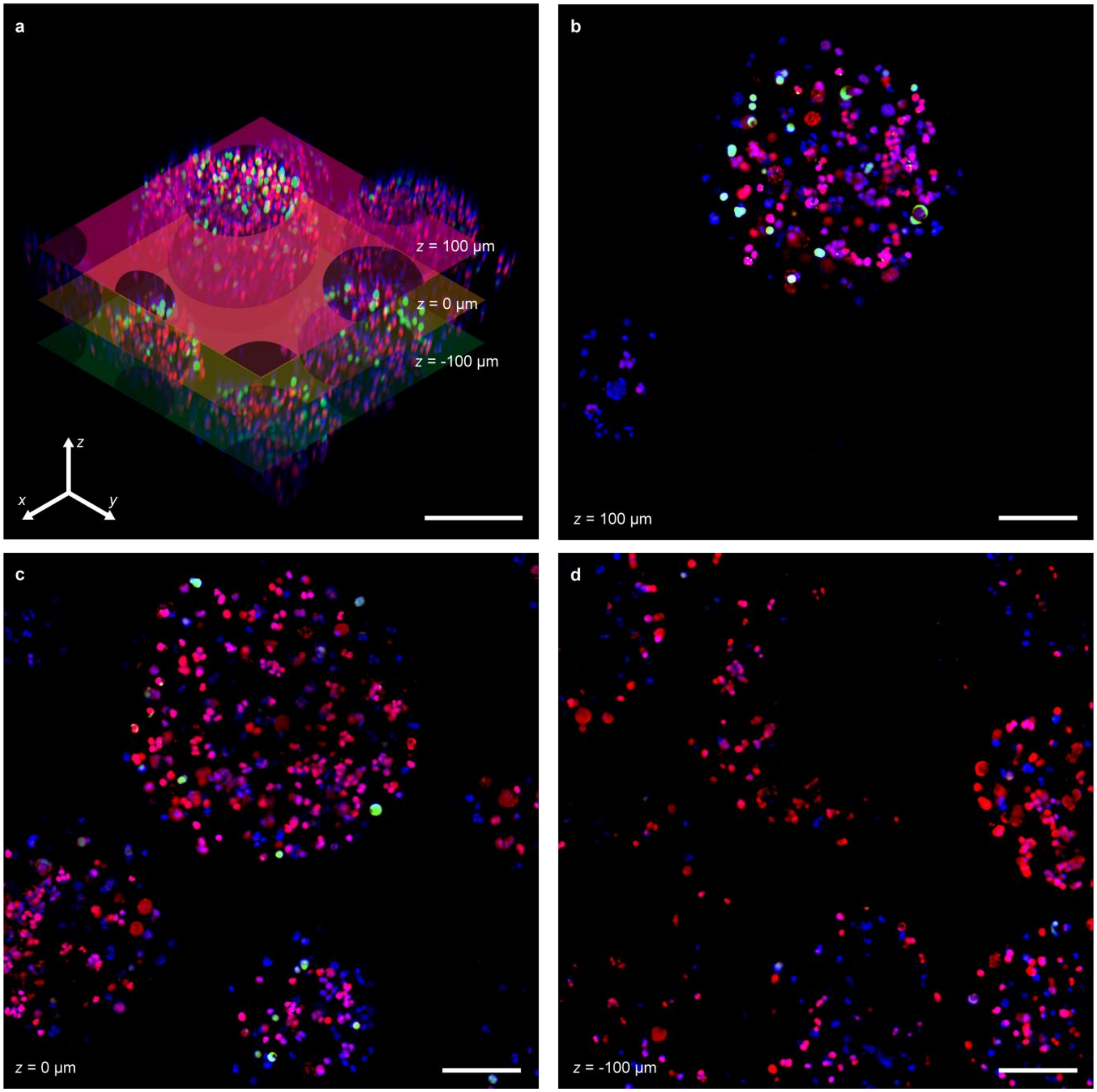


**Supplementary Figure 16. z-stack analysis of 21-day normoxia (20% $O_2$) control**; (a) 3D-reconstructed z-stacked images and the location of each presented image. (b) z=100 µm; (c) z=0 µm; (d) z=-100 µm. Scale bars are 100 µm. Green: CalceinAM (Live), Red: Ethidium homodimer (Dead), Blue: Hoechst (Nuclei).

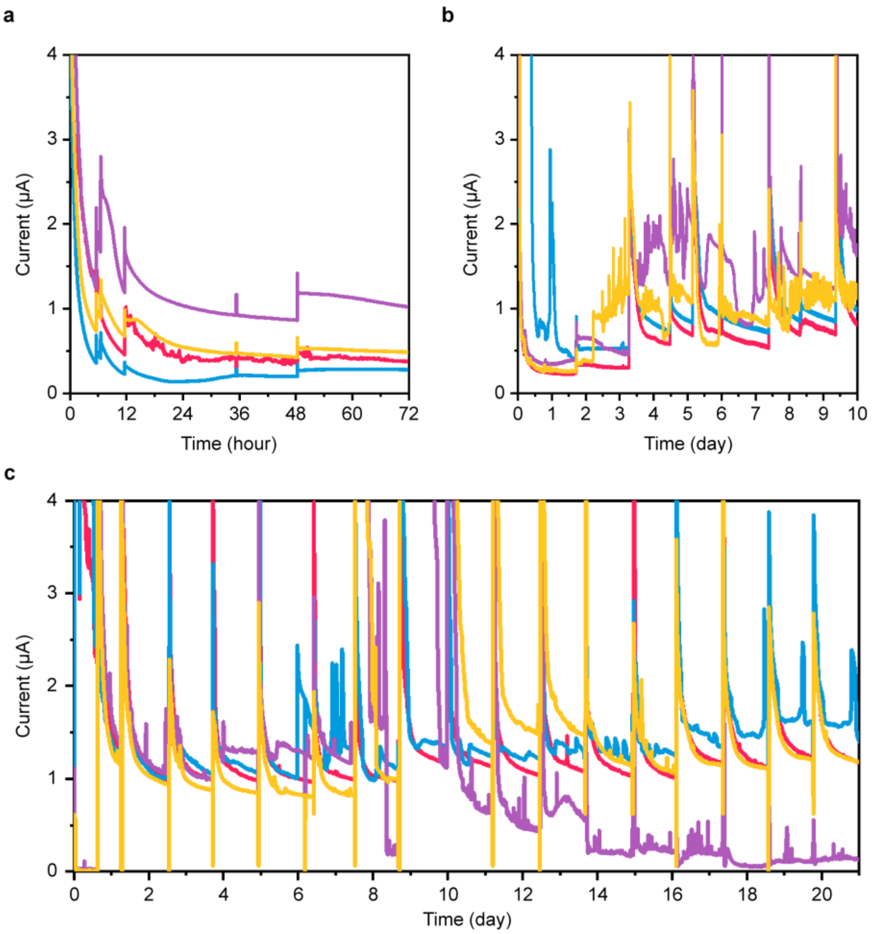


**Supplementary Figure 17. Current profiles during *in vitro* ec$O_2$ oxygenation** for (a) 3 days, (b) 10 days and (c) 21 days (n=4). Note that each spike on the profiles was caused by media exchange.

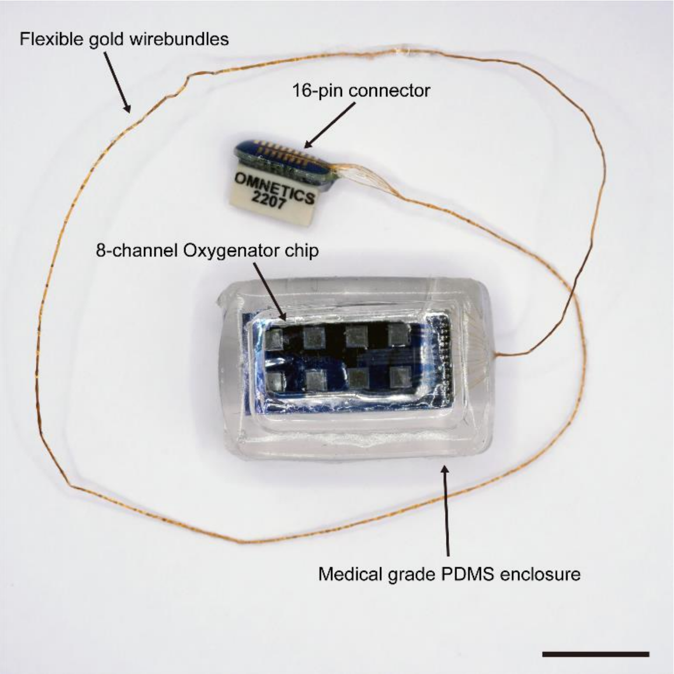


**Supplementary Figure 18. ec$O_2$ *in vivo* devices**. Note that the image was collected without a membrane; scale bar - 1 cm.

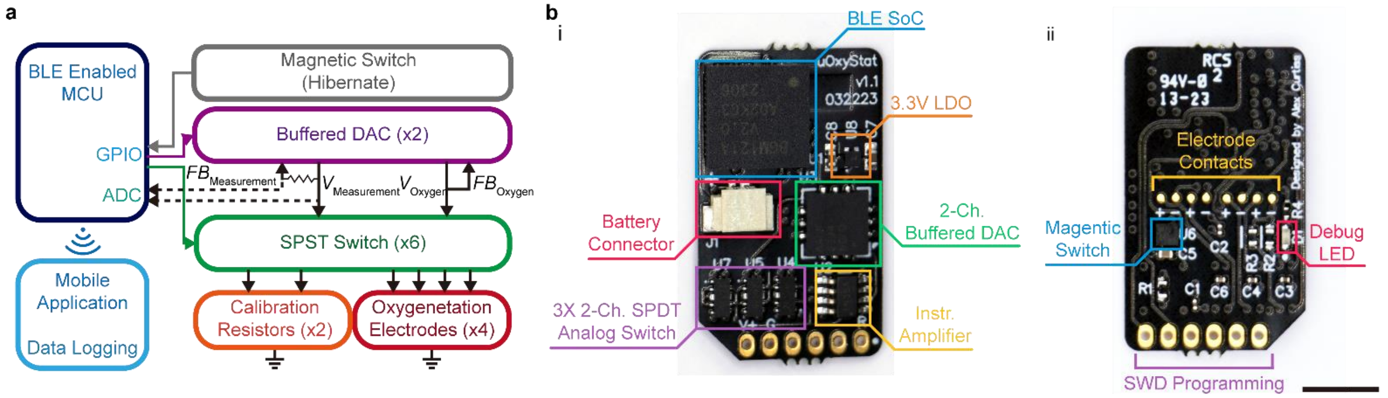


**Supplementary Figure 19. Circuit design for *in vivo* ec$O_2$.** (a) ec$O_2$ controller circuit design and (b) assembled circuits; (i) front side; (ii) backside. Scale bar is 500 µm.

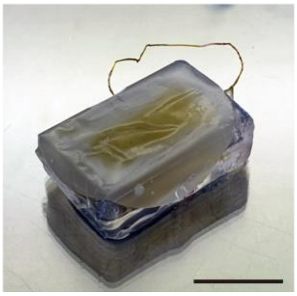

**Supplementary Figure 20. ec$O_2$ post 10 days *in vivo*.** A representative image of retrieved ec$O_2$ after 10-day. scale bar – 1 cm.

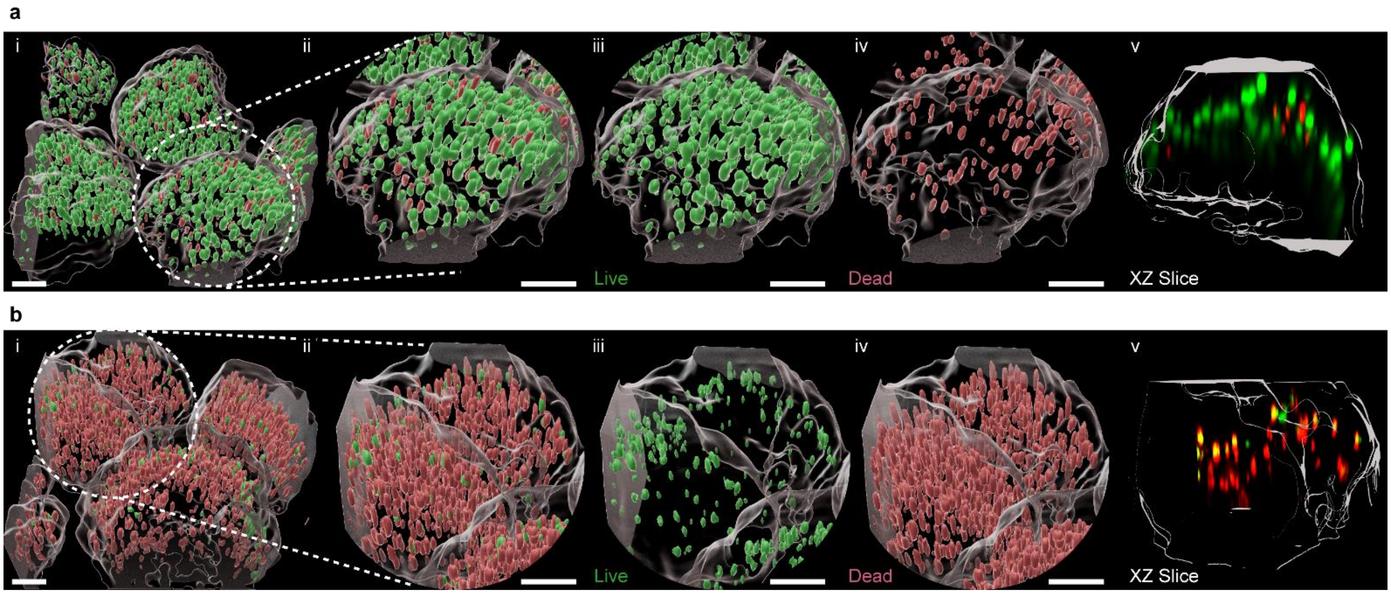


**Supplementary Figure 21. 3D-rendered fluorescence images of cell capsules post 10 days *in vivo*.** (a)

Representative image of implanted $ecO_2$ (a) with oxygenation and (b) without oxygenation. (i) 3D-rendered z-stack images. (ii-iv) expanded 3D-rendered images of capsules marked with white dashed circle. (v) Cross-sectional view in xz plane. Scale bars are 100 µm. Green: CalceinAM (Live), Red: Ethidium homodimer (Dead).

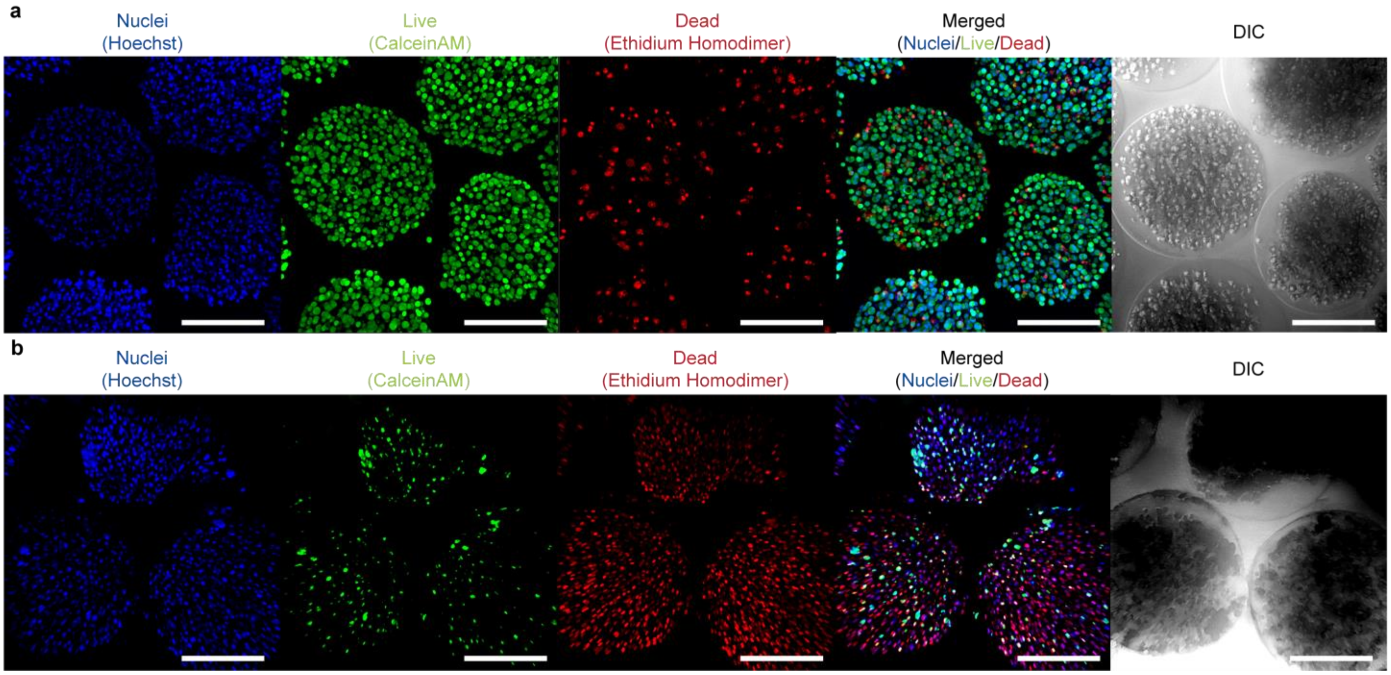


**Supplementary Figure 22. Cell viability (live/dead assay) post 10-day *in vivo*.** Representative fluorescence and optical images of capsules (a) with and (b) without oxygenation. Scale bar is 200 µm. Green: CalceinAM (Live), Red: Ethidium homodimer (Dead), Blue: Hoechst (Nuclei), DIC: differential interference contrast images (Cell capsule).

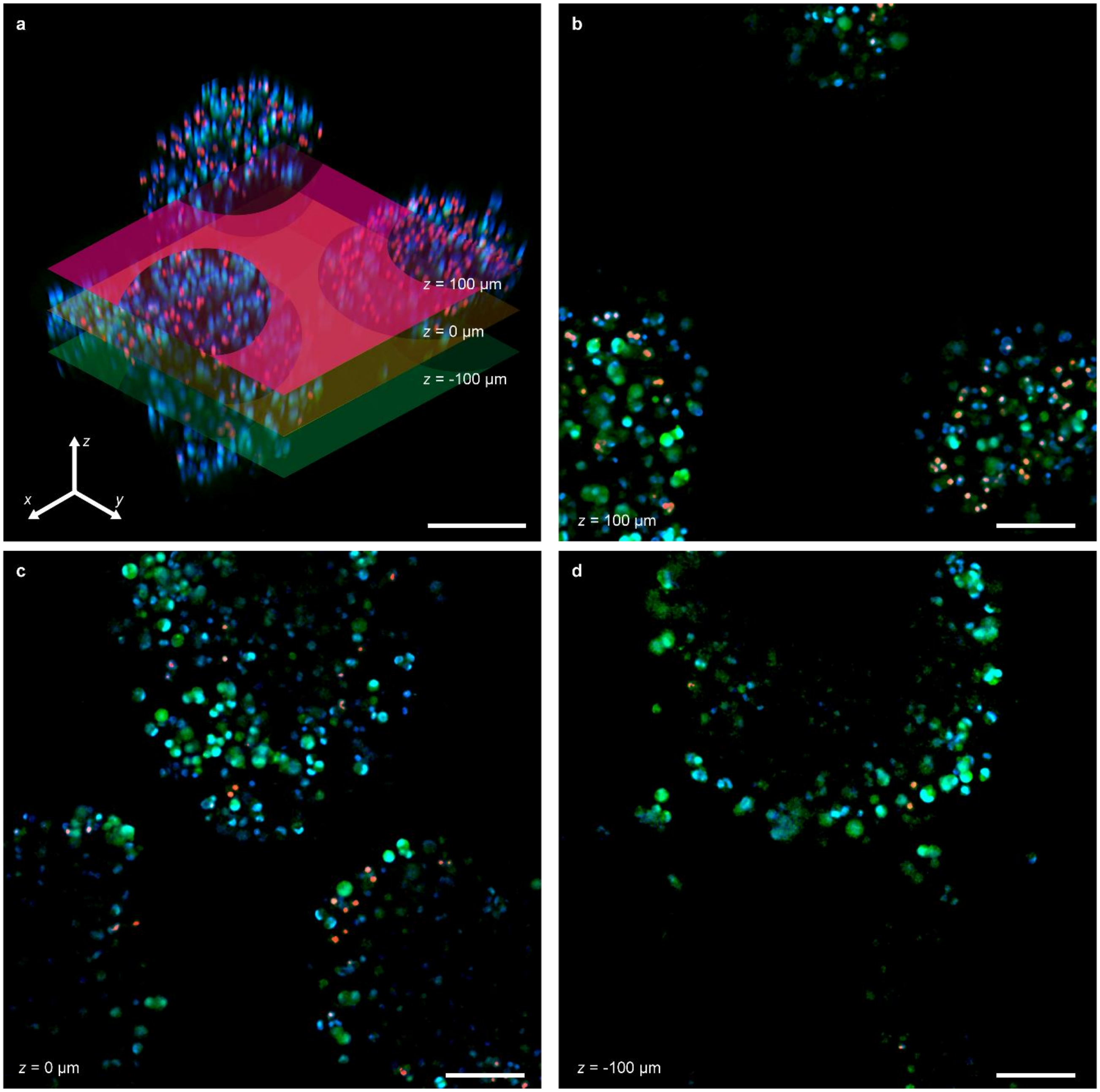


**Supplementary Figure 23. z-stack analysis of 10 days *in vivo* oxygenation**; (a) 3D-reconstructed z-stacked images and the location of each presented image. (b) z=100 μm; (c) z=0 μm; (d) z=-100 μm. Scale bars are 100 μm. WHAT ARE THE COLOR. Green: Calcein (Live), Red: B (Dead), Blue: C (Nuclei).

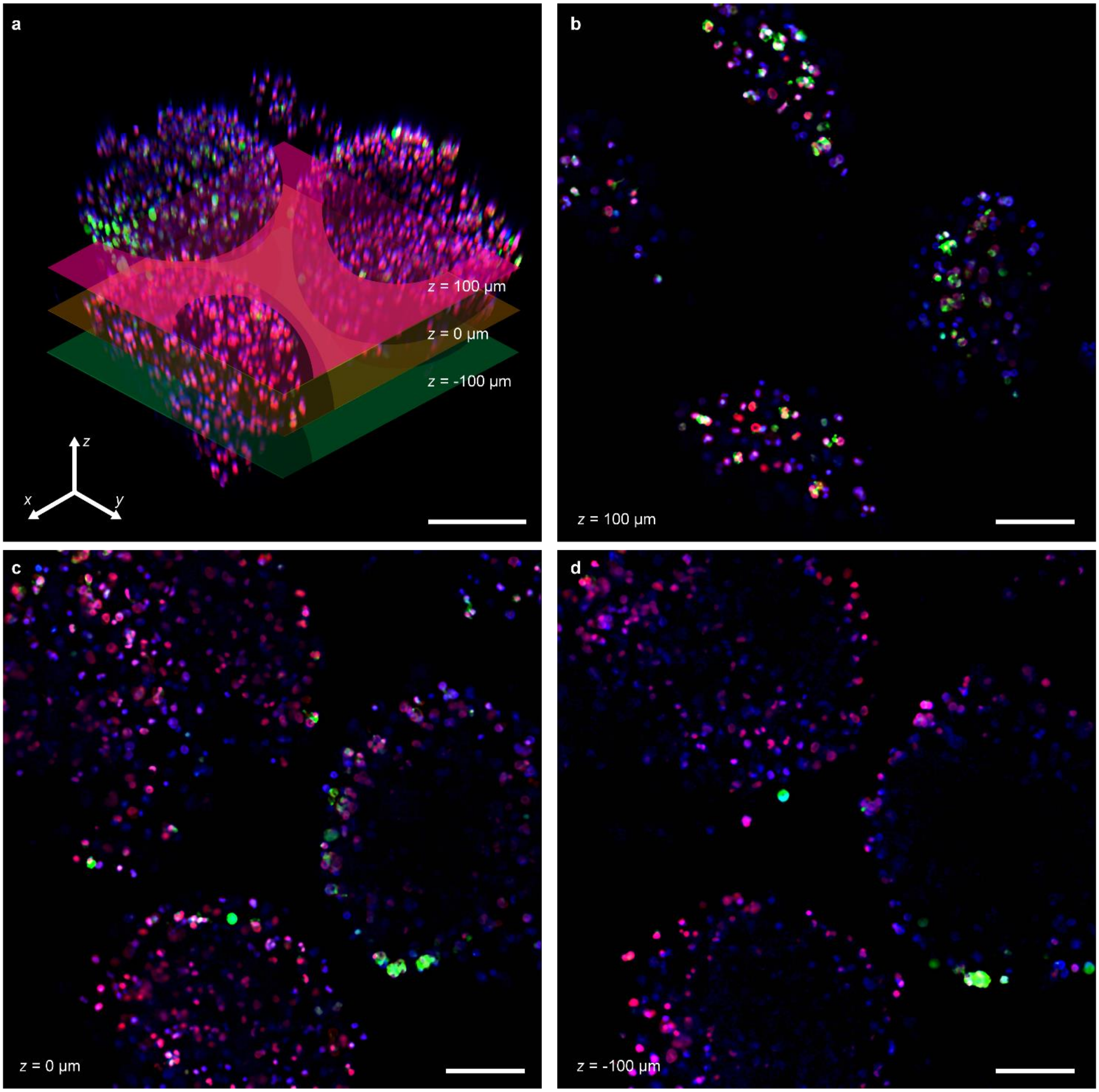


**Supplementary Figure 24. z-stack analysis of 10-day *in vivo* control**; (a) 3D-reconstructed z-stacked images and the location of each presented image. (b) z=100 μm; (c) z=0 μm; (d) z=-100 μm. Scale bars are 100 μm. WHAT ARE THE COLORS. Green: Calcein (Live), Red: B (Dead), Blue: C (Nuclei).

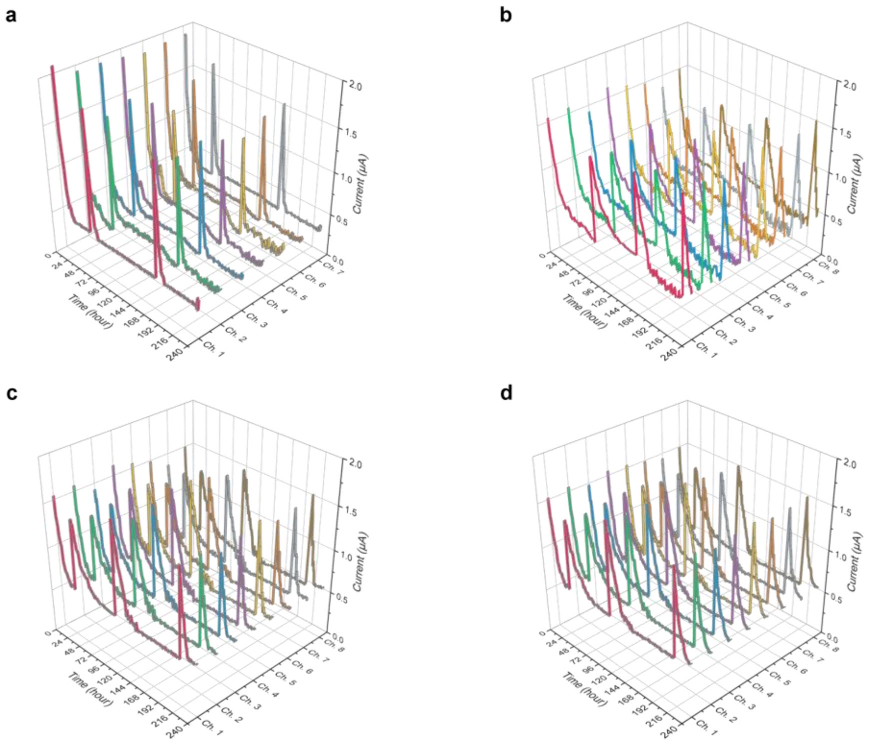


**Supplementary Figure 25. Current profiles during ecO$_2$ *in vivo*. (**a)-(d) corresponds to recorded current in active ecO$_2$ in animal 1 to 4, respectively.
